# Inhibition of human immunodeficiency virus (HIV-1) infectivity by expression of poorly or broadly neutralizing antibodies against Env in virus-producing cells

**DOI:** 10.1101/2023.09.08.556927

**Authors:** Qian Wang, Shijian Zhang, Hanh T. Nguyen, Joseph Sodroski

**Author notes:** Corresponding author: Joseph G. Sodroski, M.D. Dana-Farber Cancer Institute, 450 Brookline Avenue, CLS 1010, Boston, MA 02215. Aaron Diamond AIDS Research Center, Columbia University Vagellos College of Physicians and Surgeons, New York, NY 10032, USA.

## Abstract

The human immunodeficiency virus (HIV-1) envelope glycoprotein (Env) precursor (gp160) trimerizes, is modified by high-mannose glycans in the endoplasmic reticulum, and is transported via Golgi and non-Golgi secretory pathways to the infected cell surface. In the Golgi, gp160 is partially modified by complex carbohydrates and proteolytically cleaved to produce the mature functional Env trimer, which is preferentially incorporated into virions. Broadly neutralizing antibodies (bNAbs) generally recognize the cleaved Env trimer, whereas poorly neutralizing antibodies (pNAbs) bind the conformationally flexible gp160. We found that expression of bNAbs, pNAbs or soluble/membrane forms of the receptor, CD4, in cells producing HIV-1 all decreased viral infectivity. Four patterns of co-expressed ligand:Env were observed: 1) Ligands (CD4, soluble CD4-Ig and some pNAbs) that specifically recognize the CD4-bound Env conformation resulted in uncleaved Envs lacking complex glycans that were not incorporated into virions; 2) Other pNAbs produced Envs with some complex carbohydrates and severe defects in cleavage, which were relieved by brefeldin A treatment; 3) bNAbs that recognize gp160 as well as mature Envs resulted in Envs with some complex carbohydrates and moderate decreases in virion Env cleavage; and 4) bNAbs that preferentially recognize mature Envs produced cleaved Envs with complex glycans in cells and on virions. The low infectivity observed upon co-expression of pNAbs or CD4 could be explained by disruption of Env trafficking, reducing the level of Env and/or increasing the fraction of uncleaved Env on virions. In addition to bNAb effects on virion Env cleavage, the secreted bNAbs neutralized the co-expressed viruses.

**IMPORTANCE:** The envelope glycoprotein (Env) trimers on the human immunodeficiency virus (HIV-1) mediate virus entry into host cells. Env is synthesized in infected cells, modified by complex sugars and cleaved to form a mature, functional Env, which is incorporated into virus particles. Env elicits antibodies in infected individuals, some of which can neutralize the virus. We found that antibodies co-expressed in the virus-producing cell can disrupt Env transit to the proper compartment for cleavage and sugar modification and, in some cases, block incorporation into viruses. These studies provide insights into the processes by which Env becomes functional in the virus-producing cell and may assist attempts to interfere with these events to inhibit HIV-1 infection.

## INTRODUCTION

The human immunodeficiency virus (HIV-1) envelope glycoprotein (Env) trimer mediates virus entry into host cells and serves as a target for host neutralizing antibodies (1–5). The functional Env trimer consists of three gp120 exterior subunits and three gp41 transmembrane subunits (1,6,7). During the process of virus entry into host cells, gp120 engages the target cell receptors, CD4 and CCR5/CXCR4, and gp41 fuses the viral and cell membranes (8–21).

In infected cells, HIV-1 Env is synthesized in the rough endoplasmic reticulum (ER), where signal peptide cleavage, folding, disulfide bonding, trimerization and the addition of high-mannose glycans occur (22–25). The resulting gp160 Env precursor is transported to the Golgi apparatus, where complex sugars are added and proteolytic cleavage by host furin-like proteases produces the gp120 and gp41 subunits (26–37). The transmembrane region of gp41 anchors the Env trimer in the membrane and gp120 non-covalently associates with gp41 (38–41). The proteolytically cleaved (mature) Env trimers are transported from the Golgi compartment to the cell surface and incorporated into budding virions (42). In some Env-expressing cells, uncleaved Envs are transported to the cell surface by a secretory pathway that bypasses the Golgi and is minimally affected by brefeldin A treatment (42). Cell-surface Envs that have bypassed the Golgi compartment are not efficiently incorporated into virion particles (42).

Prior to receptor engagement, the HIV-1 Env trimer on virions can potentially sample three conformations: the pretriggered (State-1) conformation, the “open” CD4-bound (State-3) conformation, and a default intermediate (State-2) conformation (43–45). The unliganded Envs of most primary HIV-1 Envs typically exist in the pretriggered (State-1) conformation (43–46). CD4 binding initially drives Env into State 2 and then into State 3, the prehairpin intermediate conformation (43–50). In State 3, the gp41 heptad repeat (HR1) coiled coil is formed and exposed (50–53). Binding of the prehairpin intermediate to the CCR5 or CXCR4 receptor is thought to induce the formation of the six-helix bundle that mediates virus-cell membrane fusion (18–20,54–56).

The establishment of persistent infections in humans requires that HIV-1 Envs minimize the elicitation of neutralizing antibodies and resist the binding of antibodies generated during natural infection (1,2,57–62). The heavy glycan shield, surface variability and conformational lability of Env contribute to antibody evasion (63–66). During natural infection, most antibodies are elicited against forms of Env other than State 1 and therefore exhibit little or no neutralizing activity against primary strains of HIV-1 (61,62,67–69). A minority of individuals infected by HIV-1 for several years generate antibodies that broadly neutralize a variety of HIV-1 strains (70–78). Most broadly neutralizing antibodies (bNAbs) bind primary HIV-1 Envs in State 1 (43,61,62). Thus, the State-1 Envs of primary HIV-1 resist the binding of the readily elicited but poorly neutralizing antibodies (pNAbs), but are susceptible to binding and neutralization by the more rarely elicited bNAbs.

The majority of Envs synthesized in infected cells are in conformations other than State 1 and potentially can elicit pNAb responses. The gp160 Env precursor is conformationally flexible and samples many non-State-1 conformations; proteolytic maturation of gp160 in the Golgi stabilizes the State-1 conformation of the pretriggered Env trimer (79–86). Both uncleaved and cleaved Envs can bind CD4 intracellularly, adding to the population of Envs in conformations other than State 1 (87–89). As expected from the conformational diversity of the uncleaved Env, gp160 is recognized efficiently by many pNAbs (79,83,84,86,90–92). Many pNAb epitopes include gp120 elements (linear V2, V3, CD4-induced (CD4i) and Cluster A) that are exposed better upon CD4 binding (88,89,93–95). Other pNAbs recognize gp120 epitopes near the CD4-binding site, but do not bind Env trimers at the precise angle of approach required of bNAbs (96).

The isolation and characterization of monoclonal bNAbs from HIV-1-infected individuals have provided an understanding of their Env epitopes (97–101). Some bNAbs have evolved long heavy-chain complementarity-determining regions that allow access to the gp120 CD4-binding site (96,102,103). Other bNAb targets include V2 quaternary epitopes near the trimer apex, V3 glycan epitopes, hybrid gp120-gp41 epitopes and gp41 membrane-proximal external region (MPER) epitopes (104–111).

Here, we co-express pNAbs, bNAbs or CD4 with HIV-1 in virus-producing cells and find that such co-expression compromises the infectivity of the virus. The consequences of co-expression of the different ligands on Env trafficking, processing and glycosylation were examined. Distinct ligand-dependent mechanisms whereby virion Env incorporation, processing, glycosylation and function are compromised were discovered.

## RESULTS

### Co-expression of Env ligands in HIV-1-producing cells

To evaluate the effect of co-expression of Env ligands on HIV-1 infectivity, we chose several bNAbs directed against multiple Env epitopes The bNAbs included 3BNC117 against the gp120 CD4-binding site (CD4BS) (112); PGT121 against gp120 V3 glycans (106); PGT145 and PGDM1400 against gp120 V2 quaternary epitopes near the Env trimer apex (104,106,113); PGT151 and 35O22 against gp120-gp41 hybrid epitopes (107,108); and 10E8 against the gp41 MPER (110,111). We also included poorly neutralizing antibodies (pNAbs): b6 against the gp120 CD4BS (114); 19b against a gp120 V3 epitope (93); 17b and A32 against CD4-induced (CD4i) and Cluster A gp120 epitopes, respectively (88,89,94,95); and 902090 against a gp120 linear V2 epitope (115). Soluble CD4-Ig was also studied (84). Antibodies (Abs) against the Ebola virus envelope glycoprotein (2G4) (116,117), the MERS-CoV spike glycoprotein (m336) (118) and the SARS-CoV-2 spike glycoprotein (CR3022) were included as negative controls (119). The Abs produced from Expi293 cells transfected with plasmids encoding the Ab heavy and light chains were tested for the ability to neutralize HIV-1. Except for 35O22 and PGT151, the bNAbs all efficiently inhibited the infection of the primary HIV-1_AD8_ strain (Fig. 1A). The pNAbs and negative control Abs did not neutralize HIV-1_AD8_, as expected.

**FIG 1.**
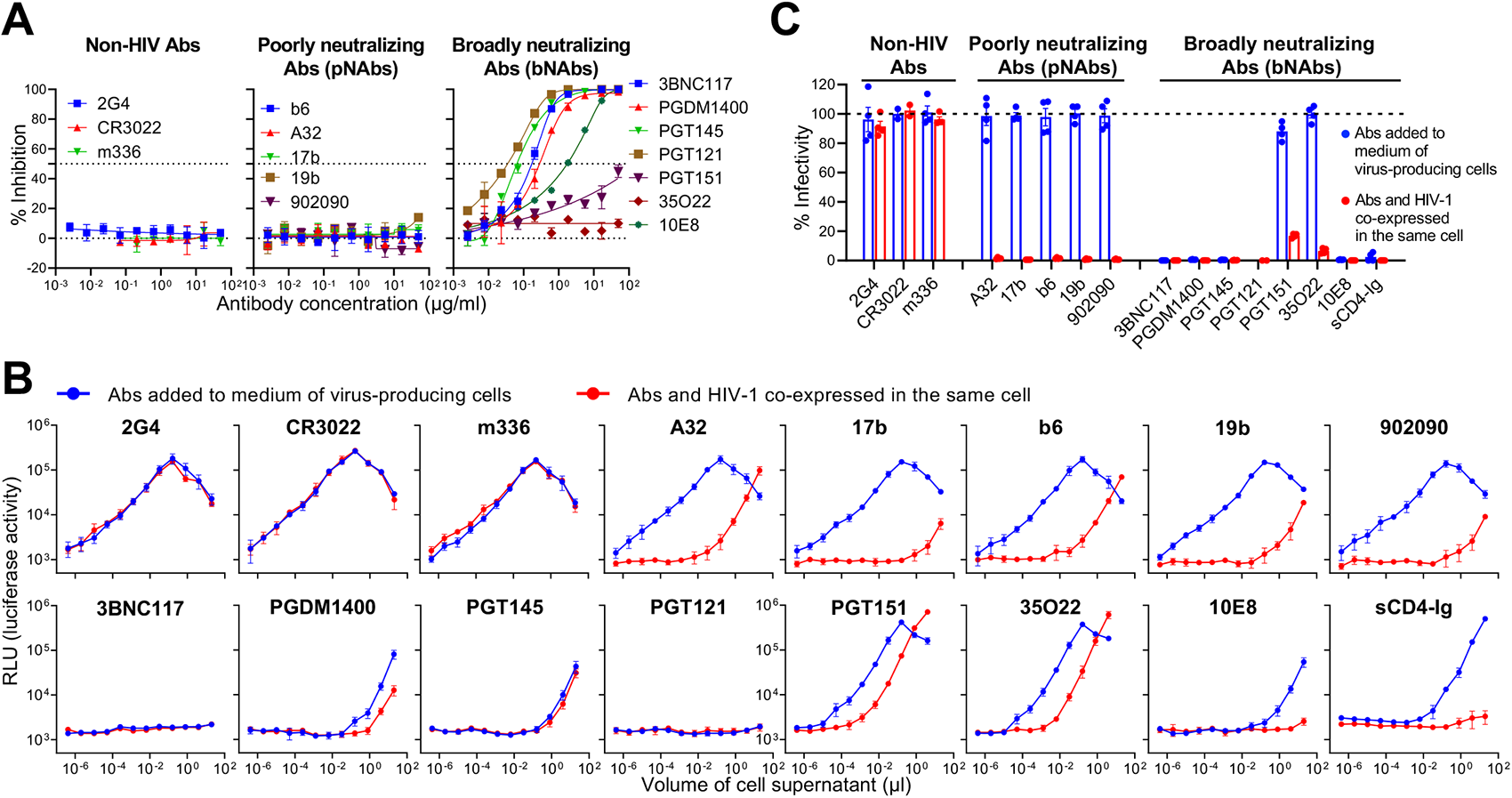
Effect of Abs on HIV-1_AD8_ infectivity. (A) Neutralization of HIV-1_AD8_ by the indicated Abs is shown. HEK293T cells were transfected with the pNL4-3-AD8 proviral plasmid to produce infectious HIV-1_AD8_, which was incubated for 1 hour at 37°C with the indicated concentrations of Abs. The virus-Ab mixtures were added to TZM-bl cells; 48 hours later, the cells were lysed and the luciferase activity (in relative light units (RLU)) in the cell lysates was measured as an indicator of virus infectivity. The % inhibition of infectivity was calculated from the observed luciferase activity relative to that of a virus not incubated with Ab. (B) The inhibition of HIV-1_AD8_ infectivity by Abs expressed in two assay formats was evaluated. In the first format (blue), HEK293T cells were transfected with the pNL4-3-AD8 plasmid. Six to eight hours later, the cell culture medium was replaced with fresh medium containing the indicated purified Abs at a concentration of 10 µg/ml. Forty-eight hours after transfection, the cell medium was collected and clarified. Different volumes of the clarified cell supernatants were added to TZM-bl cells and 48 hours later, luciferase activity in the cells was measured, as described above. In the second format (red), HEK293T cells were cotransfected with the pNL4-3-AD8 proviral plasmid and plasmids expressing the Ab heavy and light chains. Six to eight hours later, the medium was replaced with fresh medium. Forty-eight hours after transfection, the cell supernatants were harvested and clarified. The indicated volumes of the clarified supernatants were incubated with TZM-bl cells. Forty-eight hours later, luciferase activity (RLU) in the TZM-bl cells was measured, as above. The results shown represent means and standard deviations derived from triplicate samples within an experiment. (C) The % infectivity following Ab incubation, relative to that observed in the absence of antibody, is shown for experiments in which 0.4 µl of virus-containing cell supernatant was used. The results shown are representative of those obtained in two independent experiments, expressed as means and standard deviations from triplicate luciferase readings.

We then evaluated the effect of co-expressing the Env ligands on HIV-1_AD8_ infectivity. For this purpose, HEK293T cells were cotransfected with plasmids expressing the Ab heavy and light chains and the pNL4-3-AD8 proviral plasmid. Forty-eight hours later, the supernatants of the cells were titrated onto TZM-bl target cells and luciferase activity was measured. In parallel, we evaluated the effect of adding Abs to the medium of virus-producing cells. For this purpose, HEK293T cells were transfected with the pNL4-3-AD8 proviral plasmid. Six to eight hours later, purified Abs were added to the medium at 10 µg/ml final concentration. After culturing the cells for an additional 40 hours at 37°C, the cell supernatants were titrated onto TZM-bl cells and luciferase activity was measured. The luciferase activities observed for the negative control Abs 2G4, CR3022 and m336 were similar to each other in both assay formats (Fig. 1B and C). The drop in luciferase signal at larger volumes of added cell supernatants reflects viral killing of the target cells. Significant virus neutralization was observed for the 3BNC117, PGDM1400, PGT145, PGT121 and 10E8 bNAbs in both assay formats. HIV-1_AD8_ infection was minimally inhibited by the addition of the PGT151 and 35O22 bNAbs to the medium of virus-producing cells; however, modest decreases in virus infectivity were observed when these bNAbs were co-expressed with the virus. Soluble CD4-Ig inhibition of HIV-1_AD8_ infection was also more effective when co-expressed with HIV-1_AD8_ in the virus-producing cells. HIV-1_AD8_ infection was not significantly affected when the purified pNAbs (A32, 17b, b6, 19b and 902090) were added to the medium of virus-producing cells; however, when co-expressed with HIV-1_AD8_ in the virus-producing cells, all the pNAbs inhibited virus infectivity. Thus, Env ligands like sCD4-Ig and pNAbs exhibit marked increases in anti-HIV-1 inhibition when they are co-expressed with HIV-1 in the virus-producing cell. For bNAbs, several of which potently neutralized HIV-1_AD8_, more modest gains in virus-inhibitory activity were observed when the Abs were co-expressed with HIV-1 in the producing cell.

The virus-inhibitory effect of co-expressed pNAbs depended upon the amount of Ab-expressing plasmid transfected (Fig. 2A). As little as 2.34 ng of each of the heavy and light chain-expressing plasmids (compared to 150 ng of the proviral plasmid) reduced HIV-1 infectivity by approximately 70%. By contrast, no significant HIV-1_AD8_ neutralization was observed following incubation of the virus-producing cells with up to 30 µg/ml of the purified pNAbs (Fig. 2B). The antiviral effects of co-expressed Abs apparently depend on the intracellular interaction of Ab and Env.

**FIG 2.**
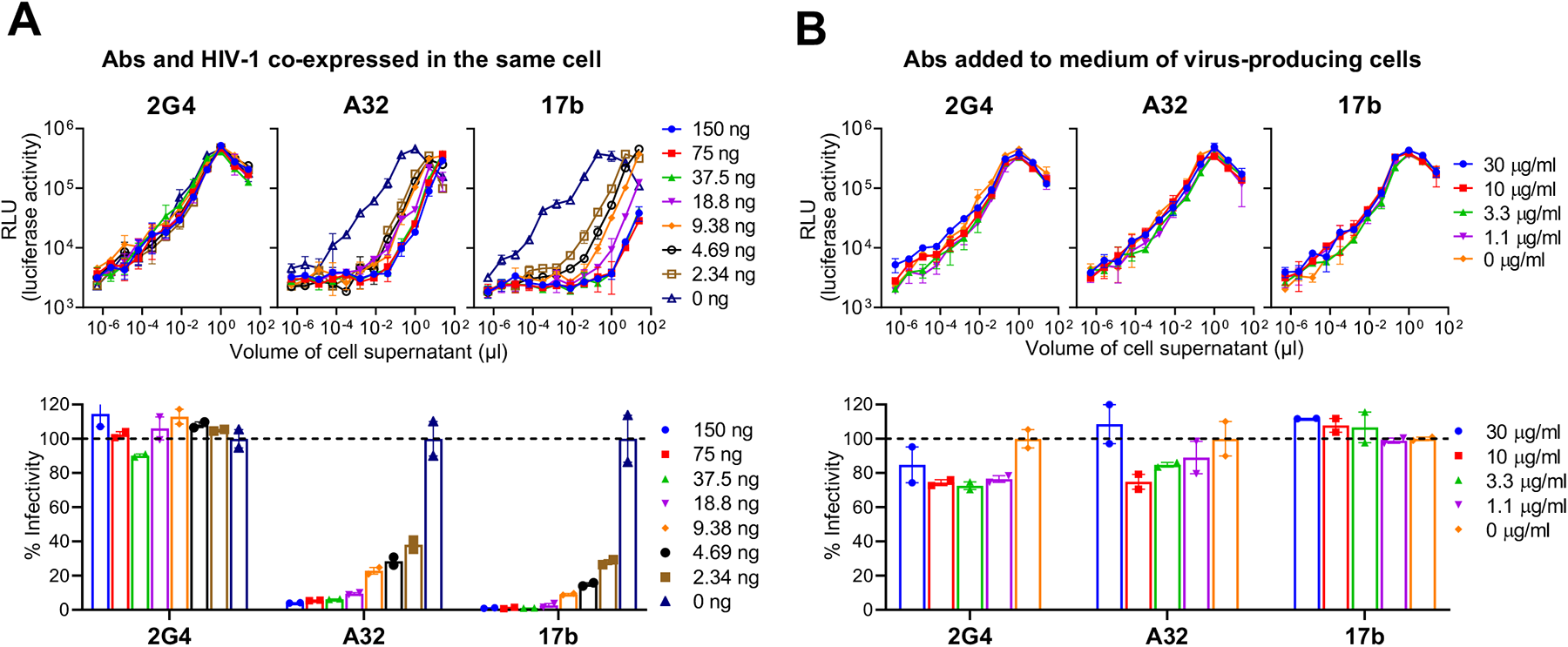
Dose-response relationships. (A) HEK293T cells were transfected with a single amount (150 ng) of the pNL4-3-AD8 proviral plasmid and the indicated total amounts of plasmids encoding the heavy and light chains of the 2G4, A32 and 17b Abs (in ng of DNA). The indicated volumes of clarified cell supernatants collected at 48 hours after transfection were incubated with TZM-bl cells. Luciferase activity (RLU) was measured in the TZM-bl cells 48 hours later. The upper panel shows the luciferase activity as a function of the volume of cell supernatant added. The lower panel shows the % infectivity using 0.4 ml cell supernatant, relative to that observed when no Ab-expressing plasmid (0 ng) was cotransfected. (B) HEK293T cells were transfected with 150 ng of the pNL4-3-AD8 plasmid. Six to eight hours later, the cell culture medium was replaced by fresh medium containing the indicated Abs at a concentration of 10 µg/ml. Clarified cell supernatants were prepared 48 hours after transfection and incubated with TZM-bl cells. Luciferase activity (RLU) in the TZM-bl cells was measured 48 hours later. The upper panel shows the luciferase activity as a function of the volume of cell supernatant added. The lower panel shows the % infectivity using 0.4 µl cell supernatant, relative to that observed in the absence of Ab (0 µg/ml). The results shown are representative of those obtained in two independent experiments, expressed as means and standard deviations from triplicate luciferase readings.

### Co-expression of pNAbs inhibits the infectivity of different HIV-1 strains

To evaluate the ability of pNAbs to inhibit the infectivity of other HIV-1 strains, we used infectious molecular clones of HIV-1_YU2_ (clade B) (120), HIV-1_CH058_ (clade B transmitted/founder) (121) and HIV-1_MJ4_ (clade C) (122). As expected, the A32 and 17b pNAbs failed to neutralize the cell-free viruses of these strains (Fig. 3A) and did not inhibit virus infectivity when the purified pNAbs were added to the medium of virus-producing cells (Fig. 3B and Fig. 3D, left panel). However, the infectivity of all three viruses was reduced when either the A32 or the 17b pNAb was co-expressed in the same cell as the provirus (Fig. 3C and Fig. 3D, right panel). Thus, pNAbs like A32 and 17b that recognize well-conserved Env epitopes can inhibit the infectivity of diverse HIV-1 when co-expressed with HIV-1 in the virus-producing cell.

**FIG 3.**
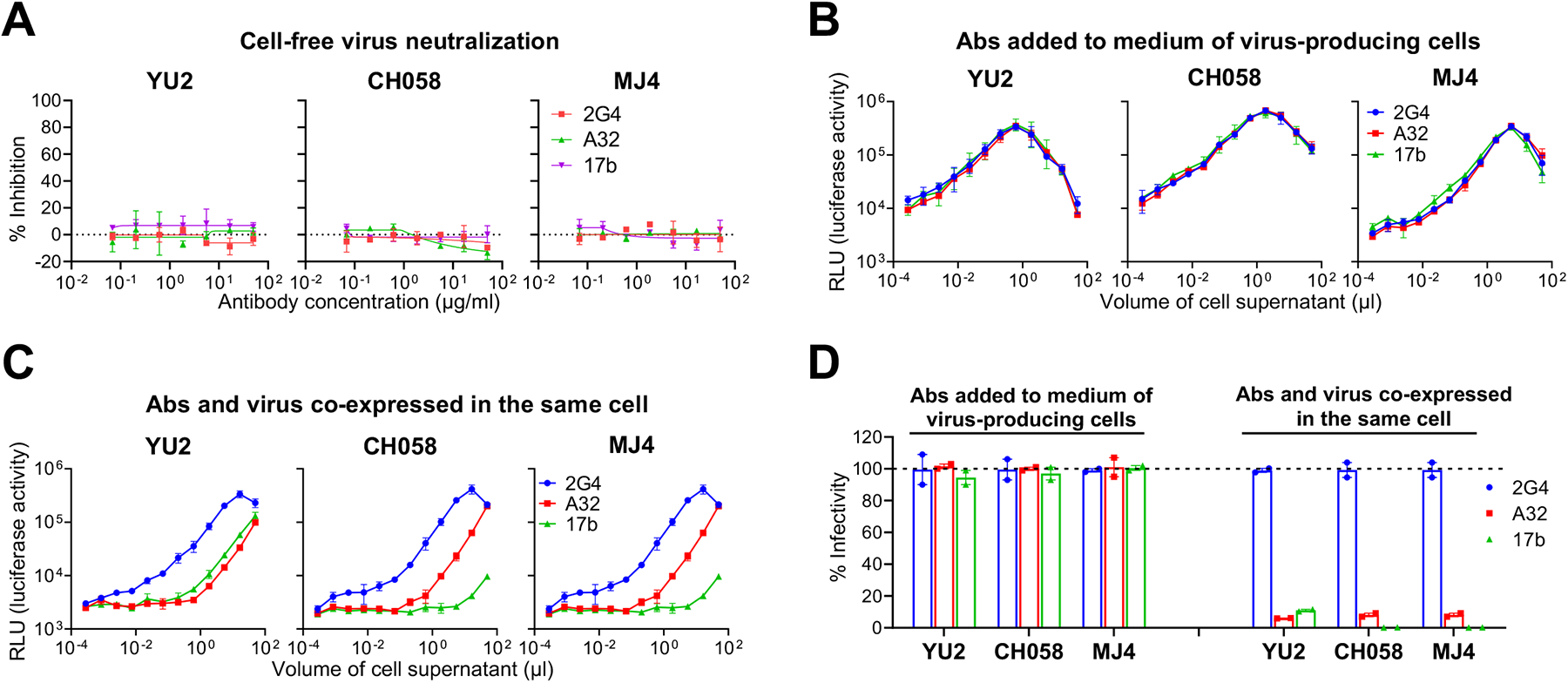
Effect of Abs on the infectivity of different HIV-1 strains. (A) Infectious clade B HIV-1_YU2_, clade B transmitted/founder HIV-1_CH058_, and clade C HIV-1_MJ4_ were produced by transfection of HEK293T cells and tested for sensitivity to neutralization by the indicated concentrations of 2G4, A32 and 17b Abs, as described in the Fig. 1A legend. The % inhibition was calculated from the observed luciferase activity in the TZM-bl target cells relative to that seen in the absence of Ab. (B) HEK293T cells were transfected with plasmids containing infectious proviruses of HIV-1_YU2_, HIV-1_CH058_ and HIV-1_MJ4_. Six to eight hours later, the cell culture medium was replaced with fresh medium containing the indicated Abs at a concentration of 10 µg/ml. Clarified cell supernatants were prepared 48 hours after transfection and incubated with TZM-bl cells. The luciferase activity (RLU) in the TZM-bl target cells following incubation with the indicated volumes of virus-containing cell supernatants is shown. (C) The indicated volumes of virus-containing supernatants from HEK293T cells cotransfected with proviral plasmids (encoding HIV-1_YU2_, HIV-1_CH058_ or HIV-1_MJ4_) and plasmids encoding the 2G4, A32 or 17b Abs were incubated with TZM-bl cells. The luciferase activity (RLU) in the TZM-bl cells was measured 48 hours later. (D) The % infectivity relative to that in the 2G4 sample is shown for the experiments in B and C at a volume of 0.4 µl cell supernatant. The results shown are expressed as means and standard deviations of triplicate samples within an experiment. The experiments were repeated with similar results.

### Effects of Env ligand co-expression on Env synthesis, processing and virion incorporation

To investigate the mechanisms underlying the effects of Env ligands on HIV-1_AD8_ infectivity, we examined the expression and processing of the Envs in cells and virions. With one exception, when purified Abs were added to the medium of virus-producing cells, the Envs in cell lysates and viral particles exhibited similar levels of expression and processing to those in control cells without added Ab (Fig. 4A, right panels). The cell and virion Envs that were exposed to the exception, sCD4-Ig, exhibited relatively low levels of gp120, likely a result of gp120 shedding (123).

**FIG 4.**
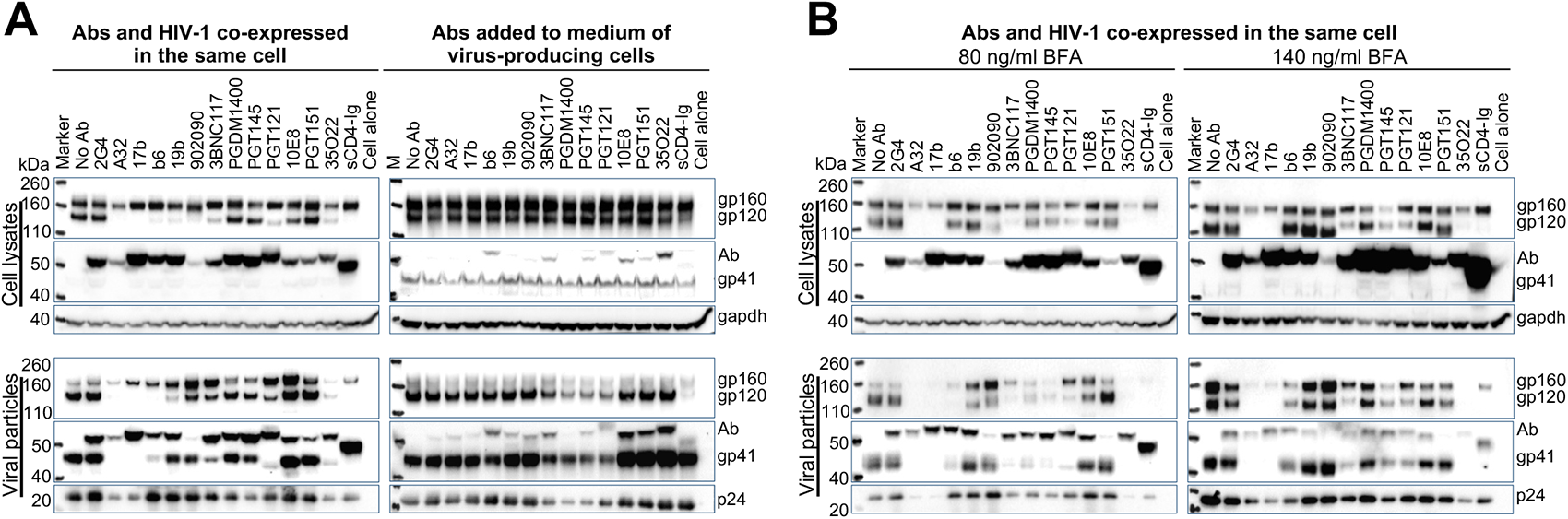
Effect of Abs on Env expression, processing and virion incorporation. (A, B) In the left panels of A and both left and right panels of B, Abs and HIV-1_AD8_ were co-expressed in HEK293T cells. In the experiments shown in the right panel of A, the medium of HEK293T cells transfected with the pNL4-3-AD8 proviral plasmid was replaced 6 hours following transfection with medium containing 10 µg/ml of the indicated Abs. In B, the transfected cells were treated with 80 ng/ml (left panels) or 140 ng/ml (right panels) brefeldin A (BFA). Cell lysates and viral particles were Western blotted for gp120 (upper panels), gp41 (middle panels), gapdh and Gag p24. In the experiments in which Abs and HIV-1_AD8_ were co-expressed, the Ab heavy chains are detected in the anti-gp41 Western blots. Note that longer exposures of the Western blots were used in the experiment in which cells were treated with 140 ng/ml BFA to allow visualization of the Envs. The results shown are typical of those obtained two independent experiments.

**FIG 5.**
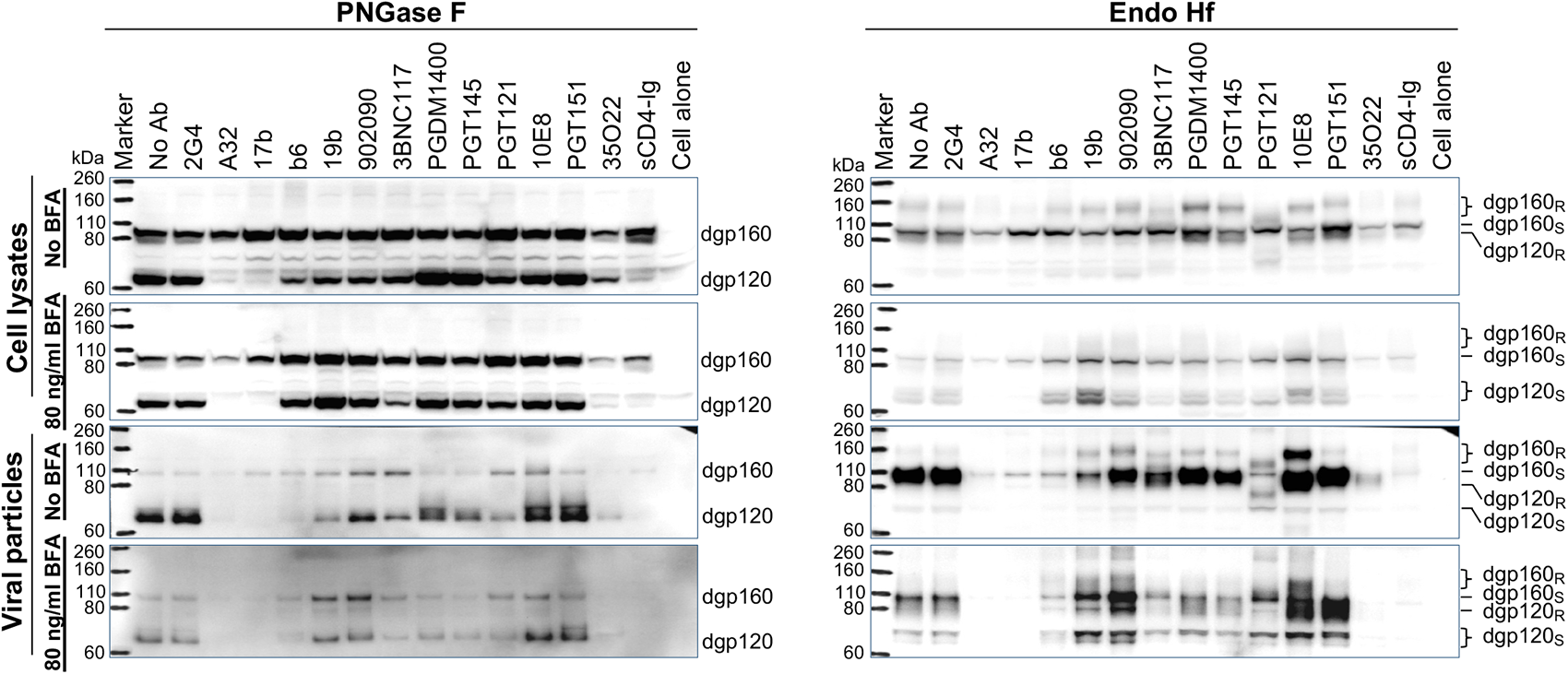
Effect of co-expressed Abs on Env processing and glycosylation. Abs and HIV-1_AD8_ were co-expressed in HEK293T cells. Cell lysates and viral particles were prepared, as described in Fig. 4 above. The cell and viral lysates were treated with PNGase F (left panels) and Endo Hf (right panels). The samples were Western blotted and the Western blots were developed with anti-gp120 Ab. The deglycosylated gp160 (dgp160) and gp120 (dgp120) proteins that were resistant (R) or sensitive (S) to Endo Hf digestion are indicated.

In the experiments in which the Abs were co-expressed with HIV-1_AD8_, we observed four patterns of Env expression and processing in cells and virions (Figs. 4 and 5; see summary in Fig. 9 below):

<u>1) sCD4-Ig and pNAbs (17b, A32) that recognize the CD4-bound Env conformation</u> – The Envs in these cells were uncleaved; the virion Envs were largely uncleaved and at low levels relative to those in control cells without Ab or co-expressing the irrelevant 2G4 Ab (Fig 4A, left panels). The vast majority of the carbohydrates on these Envs were susceptible to digestion with Endoglycosidase Hf (Endo Hf), indicating an abundance of high-mannose and/or hybrid glycans (Fig. 5). The lack of complex glycosylation and poor cleavage of these Envs were minimally affected by brefeldin A (BFA) treatment, although BFA lowered the overall level of Env expression (Figs. 4B and 5). As BFA inhibits the transport of the proteins from the endoplasmic reticulum (ER) to the Golgi (124–129), these observations are consistent with the localization of the Env-ligand complexes in the ER.
<u>2) pNAbs (b6, 19b and 902090) other than those in the above group</u> – These Envs exhibited more cleavage than those in the above group, but Env cleavage was still deficient compared with that of the control Envs (Fig. 4A, left panels). This difference in Env proteolytic processing was readily apparent when the gp120:gp160 ratios between the pNAb and control samples were compared. As is typical (42), the cleaved Envs were enriched on virus particles. Notably, the cleavage of these Envs was enhanced by treatment of the expressing cells with BFA, with cleavage efficiency increasing at a higher BFA concentration (Fig. 4B). Some complex glycans were added to these cleaved Envs either with or without BFA treatment (Fig. 5). Thus, in the absence of BFA, these Envs are at least in part transported from the ER into the Golgi. Their processing may benefit from the retrograde delivery of Golgi enzymes to the ER and/or ER-Golgi intermediate compartment (ERGIC) that occurs after BFA treatment (124–129).
<u>3) bNAbs (3BNC117, PGT121 and 10E8) that recognize gp160 as well as gp120</u> – These Envs exhibited significant decreases in the efficiency of cleavage, but the gp120 glycoproteins all contained complex carbohydrates; modest levels of total Env were incorporated into virions, although the viral gp120 levels were generally lower than those in control viruses (Fig. 4A, left panels and Fig. 5). These observations suggest that these Envs can be transported into the Golgi compartment, but are not cleaved as efficiently as the control Envs.
<u>4) bNAbs (PGDM1400, PGT145, PGT151 and 35O22) that preferentially recognize cleaved Env</u> – These Envs were cleaved and incorporated into virions efficiently, with the exception of Envs co-expressed with the 35O22 bNAb; the 35O22-associated Envs were cleaved to various degrees but were present at lower levels in the cell and virus particles (Fig. 4A, left panels). Whereas the Envs co-expressed with the PGDM1400, PGT145 and PGT151 bNAbs were minimally affected by BFA treatment of the cells, the steady-state levels of the 35O22-complexed Envs decreased upon BFA treatment. Complex glycans could be detected on all these bNAb-associated Envs (Fig. 5). With the exception of the Envs co-expressed with the 35O22 bNAb, the Envs co-expressed with these bNAbs apparently are transported through the Golgi compartment and incorporated into virus particles efficiently.

Our results suggest that, depending on the Env conformation capable of being recognized, co-expressed Env ligands inhibit Env processing and subsequent incorporation of cleaved Env into virus particles to various degrees.

### Effects of Brefeldin A (BFA) treatment on Env processing, glycosylation and virion incorporation

In the experiments shown in Figures 4 and 5 above, BFA treatment resulted in significant and, in some cases, unexpected effects on Env proteolytic processing, glycosylation and virion incorporation. Here, we present additional experimental results that provide a more complete picture of the effects of BFA in our provirus-Ab co-expression system. As in the above experiments, we utilized HEK293T cells in which the HIV-1_AD8_ provirus was expressed alone or in combination with Env-reactive Abs. The cells were treated with different concentrations of BFA for approximately 40 hours, after which lysates of cells and viral particles were used to analyze the Env glycoproteins. In the absence of co-expressed Ab, the effects of BFA treatment were concentration dependent (Fig. 6A). At low concentrations (∼80 ng/ml), BFA treatment resulted in mild decreases in the proteolytic processing of Env in cell lysates; this decrease in Env processing was less pronounced in virions, where cleaved Env is typically enriched (42). At higher concentrations (>120 ng/ml), BFA treatment resulted in substantial increases in the migration of both gp160 and gp120 in cells and virions; this increase in migration results from altered glycosylation, as indicated by the analysis of the Envs after PNGase F digestion (Fig. 6A). Also, progressive increases in the levels of Env in the cell lysates and decreases in virion-associated Env were observed as BFA concentration increased (Fig. 6A). On the other hand, the release of viral particles into the cell medium was not significantly affected by BFA treatment. Apparently, treatment with higher concentrations of BFA for approximately two days decreased the efficiency of Env incorporation into viral particles.

**FIG 6.**
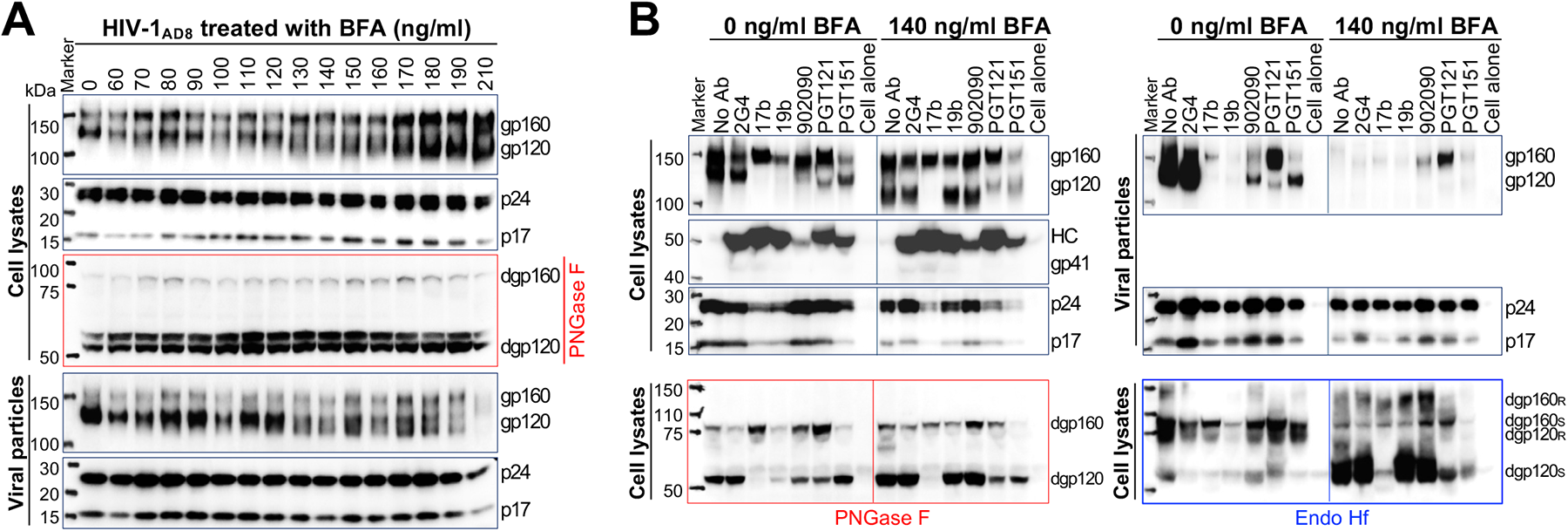
Effect of brefeldin A (BFA) on Env processing, glycosylation and virion incorporation. (A) HEK293T cells were transfected with the pNL4-3-AD8 proviral plasmid. Six hours after transfection, the cells were transferred into fresh medium containing the indicated concentrations of BFA. Forty-eight hours after transfection, cell lysates and viral particles were prepared. Portions of the cell lysates were treated with PNGase F. Cell lysates and viral particles were Western blotted and blots were developed with anti-gp120 Abs and anti-Gag Abs. The results shown are typical of those obtained in two independent experiments. (B) HEK293T cells were cotransfected with the pNL4-3-AD8 proviral plasmid and plasmids expressing the heavy and light chains of the indicated Abs. Six hours after transfection, the cells were transferred into fresh medium with the indicated concentrations of BFA. Forty-eight hours after transfection, cell lysates and viral particles were prepared for Western blotting. In the lower panels, cell lysates were treated with PNGase F and Endo Hf for 1 hour before Western blotting. The deglycosylated gp160 and gp120 are labeled as dgp160 and dgp120, respectively. Proteins that were resistant (R) or sensitive (S) to Endo Hf digestion are indicated. The results shown are typical of those obtained in two independent experiments.

The concentration dependence of the observed effects of BFA treatment on Env processing, glycosylation and virion incorporation provides a rationale for examining the effects of BFA treatment at 80-and 140-ng/ml (see Fig. 4B). As the effects of 80 ng/ml BFA on Env processing and glycosylation are presented in greater detail in Figure 5, we provide additional data on the effects of 140 ng/ml BFA in Figure 6B. Comparison of the results at the two BFA concentrations reveals interesting similarities and differences in Env processing, glycosylation and virion incorporation. In the absence of BFA, nearly all of gp120 and some gp160 was modified by complex carbohydrates; cleaved Envs and a smaller amount of gp160 with complex glycans were enriched in virions (Fig. 5), as previously seen (42,92). Treatment with 80 ng/ml BFA resulted in decreases in the levels of complex carbohydrates added to both gp160 and gp120 in cells and virions (Fig. 5). The BFA-induced increases in Env cleavage associated with co-expression of the b6, 19b and 902090 pNAbs resulted in the unusual appearance of Endo Hf-sensitive gp120, indicative of extensive modification by high-mannose and/or hybrid glycans.

In Figure 6B, untreated and 140 ng/ml BFA-treated HEK293T cells co-expressing HIV-1_AD8_ and representatives of the different Ab groups were compared. As seen at lower BFA concentrations, dramatic increases in the proteolytic processing of the Envs co-expressed with the 19b and 902090 pNAbs were observed with 140 ng/ml BFA. Equally impressive decreases in the virion incorporation of all the Envs, regardless of co-expressed Abs, were observed after two days of treatment with 140 ng/ml BFA. At this BFA concentration, nearly all of the glycans on gp120 were Endo Hf-sensitive and therefore of the high-mannose/hybrid type, regardless of the co-expressed Ab. Thus, further reductions in the degree of complex carbohydrate modification of cleaved Envs seen at 80 ng/ml BFA were observed at 140 ng/ml BFA. In contrast to this effect and the results seen at 80 ng/ml BFA, more complex (Endo Hf-resistant) carbohydrates were added to gp160 in the presence of 140 ng/ml BFA. One exception was the Env co-expressed with PGT121; recognition of a gp120 outer domain V3 glycan epitope by this bNAb (106) apparently influences the modification of the co-expressed Env by the host glycosylation machinery (Figs. 5 and 6B). In summary, at these concentrations and treatment durations, BFA can increase the cleavage of Envs co-expressed with certain pNAbs, differentially alter the content of complex glycans on gp160 and gp120, and generally reduce Env incorporation into virions. These observations are integrated into a consistent model in the Discussion (see Fig. 9 below).

### Effects of Ab co-expression on cell-surface Env

To examine the effects of Ab co-expression on cell-surface Env, HEK293T cells expressing HIV-1_AD8_ and Abs were biotinylated. Envs on the cell surface were analyzed in parallel with Envs in the total cell lysates and viral particles (Fig. 7). Consistent with our results above (Figs. 4A and 5), by 24 hours after transfection, significant decreases in the processing of cell-lysate Envs were associated with co-expression of all the Abs except the PGT145 and PGDM1400 bNAbs. Decreases in the levels of virion gp120 accompanied co-expression of the pNAbs. With the exception of PGT145 and PGDM1400, co-expression of the bNAbs led to decreases in the level and/or ratio of cleaved/uncleaved Env on virions. For most of the Abs, the pattern of Envs on the cell surface paralleled that in the total cell lysates. Thus, as previously seen (42), both uncleaved and cleaved Envs can be efficiently transported to the surface of expressing cells.

**FIG 7.**
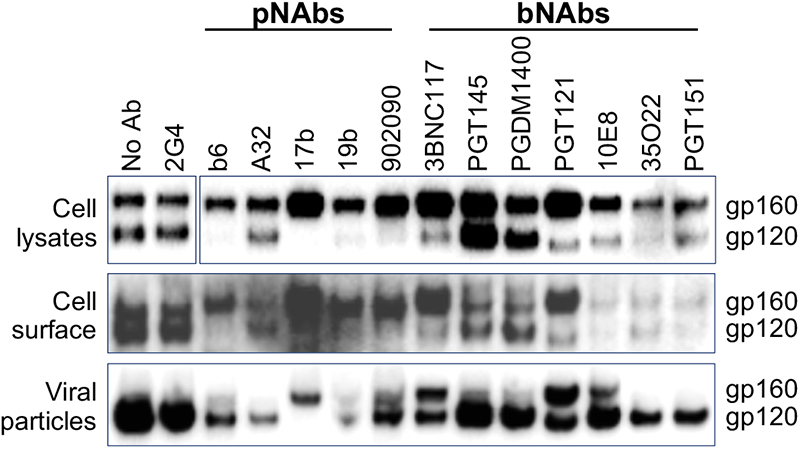
Cell-surface expression of HIV-1_AD8_ Envs co-expressed with Abs. HEK293T cells were transfected with the pNL4-3-AD8 proviral plasmid and with plasmids expressing the indicated Abs. Approximately 24 hours after transfection, Envs in lysates of cells and viral particles were analyzed by Western blotting with a goat anti-gp120 polyclonal Ab, as described in Materials and Methods. A fraction of the transfected cells was used for biotinylation of cell-surface proteins. After washing with 1x glycine in PBS, the cells were lysed. Clarified cell lysates were incubated with NeutrAvidin-Agarose resin and the captured proteins were Western blotted with a goat anti-gp120 polyclonal Ab. The results of a typical experiment are shown.

### Effect of CD4 co-expression on Env processing and virion incorporation

The results in Figs. 4A and 5 showed that co-expression of soluble two-domain CD4 (sCD4-Ig) with Env reduced Env processing and virion incorporation. We compared the effect of co-expression of full-length, membrane-anchored CD4 with that of sCD4-Ig on Env processing. Env cleavage in cell lysates was dramatically reduced by co-expression of either form of CD4 (Fig. 8A). We examined the effect of co-expression of full-length, membrane-anchored CD4 on HIV-1_AD8_ Env processing, cell surface expression and incorporation into virus particles. In the experiment shown in Figure 8B, the HIV-1_AD8_ Env and CD4 were expressed individually or together in HEK293T cells. Co-expression of CD4 completely inhibited the processing of Env in the cell lysates and on the cell surface (Fig. 8B). The cell-surface gp160 was precipitated by the OKT4 anti-CD4 antibody in cells co-expressing CD4, indicating that Env-CD4 complexes trafficked to the cell surface. The cell-surface gp160 was sensitive to Endo Hf digestion and therefore was not modified by complex glycans.

**FIG 8.**
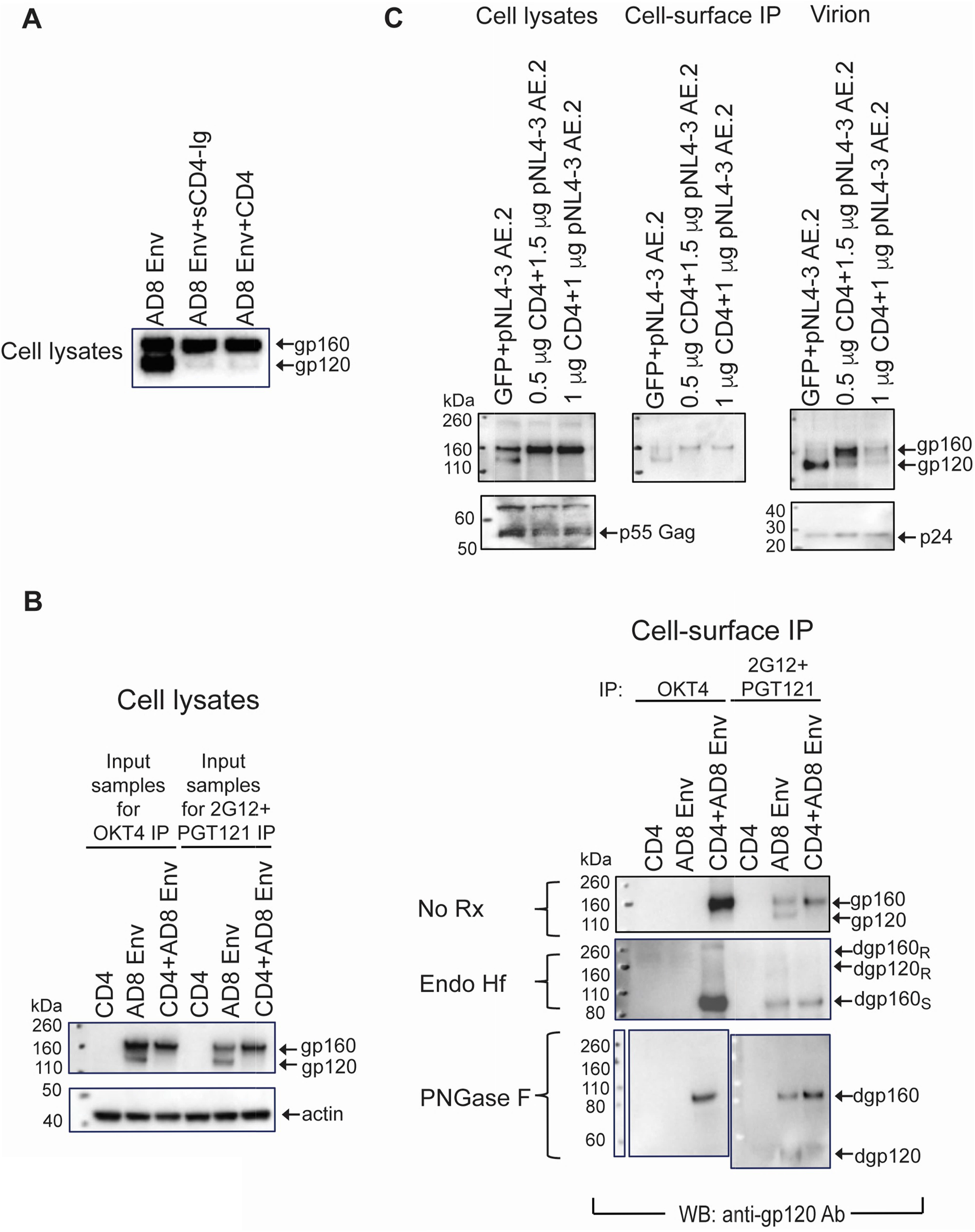
Effect of co-expression of CD4 on Env processing, glycosylation and virion incorporation. HEK293T cells were transfected with pNL4-3-AD8 (A, B) or pNL4-3-AE.2 (C) proviral plasmids, either with or without plasmids expressing sCD4-Ig or full-length human CD4. Forty-eight hours after transfection, Envs in the cell lysates and virus particles were analyzed by Western blotting with an anti-gp120 Ab. In B and C, cell-surface Envs were precipitated with a mixture of 2G12 and PGT121 bNAbs against gp120. In B, cell-surface Env:CD4 complexes were precipitated with the OKT4 Ab against CD4. The cell-surface immunoprecipitates in B were treated with Endo Hf and PNGase F. The cell-surface immunoprecipitates in B and C were Western blotted with an anti-gp120 Ab. Deglycosylated gp160 (dgp160) and gp120 (dgp120) that were resistant (R) or sensitive (S) to Endo Hf digestion are indicated. The results shown are typical of those obtained in two independent experiments.

**FIG 9.**
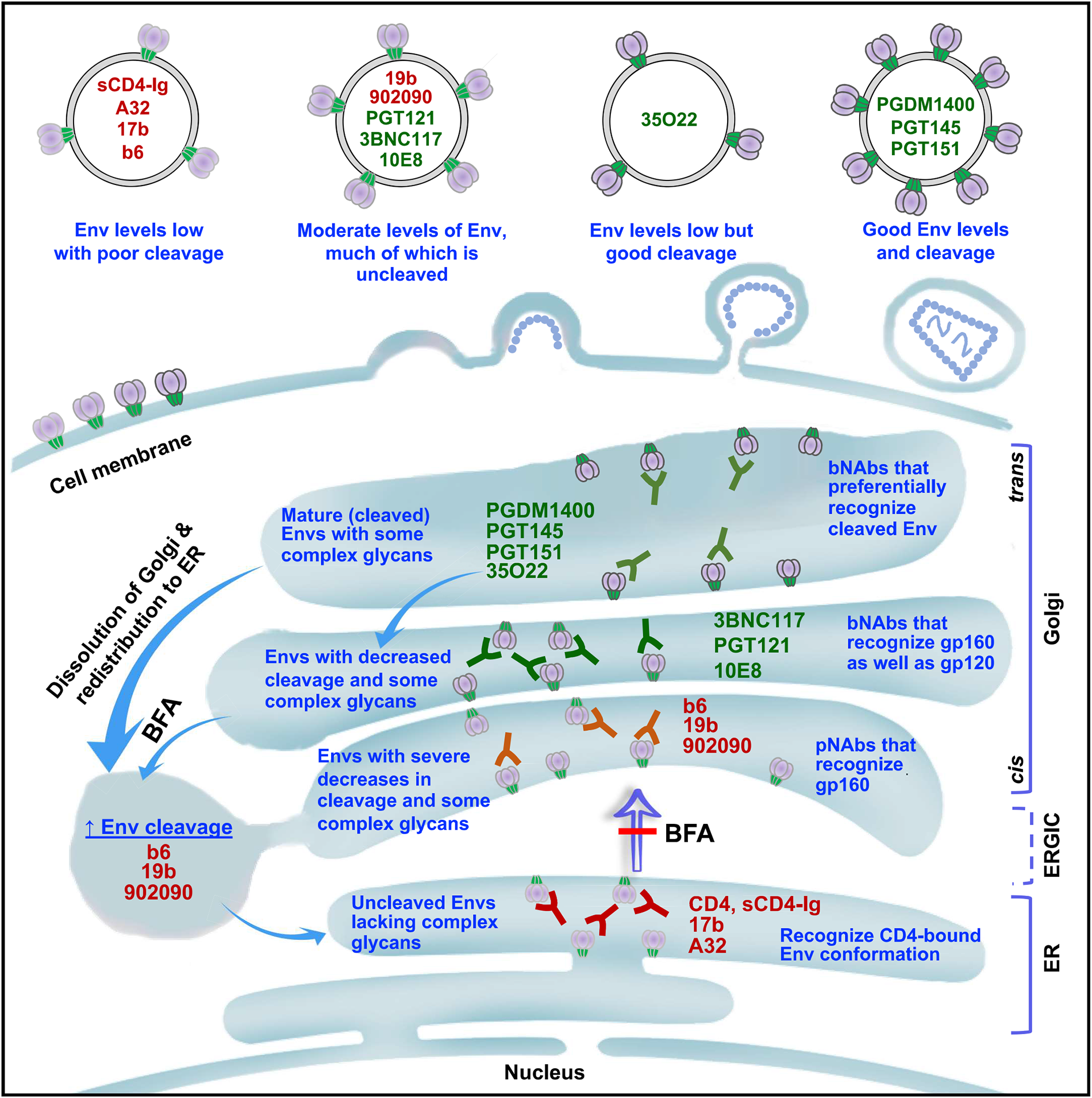
A model summarizing the effects of co-expressed Abs on Env. HIV-1 Envs are synthesized in the endoplasmic reticulum (ER) and are transported through the classical secretory pathway to the cell surface, from which the Envs are incorporated into virions. From the ER, Envs move through the ER-Golgi intermediate compartment (ERGIC) and the *cis*, medial and *trans* Golgi to the cell surface and virions. In the distal Golgi, complex carbohydrates are added to Env and proteolytic cleavage occurs, producing the mature Env that is incorporated into virions. Co-expressed Env ligands (CD4 and Abs) recognize Envs at different stages of folding and maturation, as indicated to the right of the ligand names (bNAbs are colored green, pNAbs and CD4 red). The Envs that result from interaction with the Abs or CD4 are indicated to the left of the ligand names. The consequences of ligand co-expression on the level and processing of Env on virion particles are illustrated. Brefeldin A (BFA) inhibits anterograde transport from the ER. BFA treatment also results in dissolution of more distal Golgi components, which redistribute to the ER and ERGIC. Thus, upon treatment with BFA, Golgi enzymes involved in Env cleavage and glycosylation can act upon some Envs trapped in earlier compartments of the secretory pathway (e.g., as a result of interaction with the b6, 19b and 902090 pNAbs).

To evaluate the effect of CD4 co-expression on Env incorporation into virus particles, we used the pNL4-3-AE.2 proviral clone. This HIV-1 provirus expresses the AE.2 Env, an AD8 Env variant that is processed more efficiently than the HIV-1_AD8_ Env (130). HEK293T cells were transfected with pNL4-3-AE.2 alone or with two amounts of a CD4-expressing plasmid. Cotransfection of either amount of the CD4 expressor plasmid resulted in decreased Env processing on the cell surface and on virions (Fig. 8C). At the higher amount of the transfected CD4 plasmid, the level of Env on the virion decreased, even though the level of cell-surface gp160 did not. Thus, even at low levels, co-expression of full-length CD4 resulted in decreased Env processing. At high levels of CD4 expression, the incorporation of Env into virions was diminished. These inhibitory effects on Env cleavage and virion incorporation are similar to those seen above for co-expression of sCD4-Ig with HIV-1_AD8_.

## DISCUSSION

In this work, we measured the effect of anti-Env Abs on the infectivity of viruses produced from cells transfected with infectious HIV-1 proviral clones. The Abs were tested in two formats. In one format, a fixed concentration of each purified Ab was added to the medium of provirus-expressing cells. In this format, bNAbs decreased the infectivity of the produced virus to varying degrees, whereas pNAbs did not. With the exception of sCD4-Ig, which induced shedding of gp120, the Abs did not detectably affect the integrity of the virion Envs in this format. Only some bNAbs against the gp120 CD4BS (e.g., b12 but not VRC01) and gp41 MPER (e.g., 2F5 and 4E10) have been reported to induce gp120 shedding; however, this Ab-induced gp120 shedding is much less efficient and slower than shedding induced by sCD4, requiring several hours of incubation (131,132).

In a second format, when the Abs were co-expressed in the same cell as the HIV-1 provirus, all of the anti-Env Abs decreased the infectivity of the produced virus to some extent. The potency of this inhibition was not dependent on the ability of the Ab to neutralize the virus, as some co-expressed pNAbs inhibited virus infectivity better than some bNAbs. However, the inhibitory activity of the co-expressed Ab was always as great or greater than the virus-neutralizing activity of the same Ab. This observation suggests that the inhibition of virus infectivity observed with Ab co-expression is the sum of extracellular neutralization and any intracellular events that further compromise Env integrity or function. Therefore, bNAb neutralization potency is one factor that contributes to the rank order of potency of co-expressed Abs. Another factor appears to be the stage of Env synthesis, processing and transport at which the Ab engages Env (Fig. 9). Abs that can bind at an earlier stage potentially block processes that are critical for virus infectivity (Env folding, proteolytic cleavage, transport through the classical secretory pathway leading to virion incorporation (42)). A significant part of the antiviral activity of the co-expressed pNAbs, which engage immature Envs, can be attributed to disruption of these processes. On the other hand, bNAbs such as PGDM1400, PGT145, PGT151 and 35O22 that preferentially recognize cleaved Env cannot engage Env until late in its secretion through the Golgi; thus, when co-expressed with HIV-1, the virus-inhibitory activity of these Abs was only marginally better than their extracellular neutralizing activity. We also showed that the antiviral activity of the co-expressed Abs increased at higher concentrations; nonetheless, at saturation, the co-expressed 17b pNAb was more potent than the A32 pNAb. Thus, other factors like the level of epitope exposure, the Ab affinity, and the consequences of Ab binding to Env folding and processing conceivably contribute to the antiviral potency of a given co-expressed Ab.

Several consequences of engagement of Env with the co-expressed Ab were observed that likely contribute to the decrease in infectivity of the virus produced from the cells (Fig. 9). The pNAbs 17b and A32 preferentially recognize CD4-bound Env conformations and, like sCD4-Ig and CD4 itself, strongly inhibited Env proteolytic processing and modification by complex glycans. These ligands likely arrest Env transport from the ER, denying access to the Golgi where proteolytic cleavage and complex carbohydrate modification occur (24,30,32–34). In addition, sCD4-Ig and CD4 can induce open Env conformations that are functionally labile (123,133). Other pNAbs like b6, 19b and 902090 also promote significant reductions in Env cleavage, although not as completely as the Env ligands above. Some bNAbs like 3BNC117, PGT121 and 10E8 recognize gp160 as well as gp120 and decrease the efficiency of Env cleavage. As mentioned above, the bNAbs that preferentially recognize cleaved Env (PGDM1400, PGT145 and PGT151) bind too late to intervene in the cleavage event.

The effects of the co-expressed Abs on Env folding, processing or transport ultimately impact the level and maturity of Env on virions (Fig. 9). Except for the bNAbs PGDM1400, PGT145 and PGT151, where virions from co-expressing cells exhibited near-normal levels of cleaved Envs, virions produced from cells co-expressing anti-Env Abs exhibited lower levels of Env, a higher percentage of which was uncleaved. Should this uncleaved Env be mixed with cleaved Env protomers, the resultant Env trimers will be functionally compromised (134). Thus, the Abs that bind gp160 and inhibit its subsequent processing can significantly impair virus infectivity.

Evaluating Env resistance to Endo Hf digestion, indicative of the acquisition of complex carbohydrates, in conjunction with the effects of BFA treatment can provide information on Env progression through the secretory pathway. BFA slows the anterograde transport of proteins from the ER to the ER-Golgi intermediate compartment (ERGIC) and Golgi apparatus (135). In addition, BFA induces a rapid disassembly of Golgi complexes and a relocation of Golgi enzymes to the ER (124–129). We found that, at low BFA concentrations, complex glycan modification of both gp160 and gp120 was diminished compared with untreated controls. At higher BFA concentrations, while gp120 exhibited even higher levels of high-mannose/hybrid glycans, gp160 demonstrated an increase in the addition of complex glycans. Moreover, the poorly processed Envs co-expressed with the b6, 19b and 902090 pNAbs demonstrated impressive increases in cleavage following BFA treatment. These properties of Env observed after BFA treatment can be explained by the relocation of Golgi enzymes (furin-like proteases, glycosylation enzymes) to the ER and ERGIC (124–129). Thus, Envs made in cells co-expressing the b6, 19b and 902090 pNAbs, which in the absence of BFA appear to be located in the ERGIC and/or early (cis) Golgi, could be exposed to higher amounts of furin-like proteases retrotransported from late (trans) Golgi compartments after BFA treatment. Likewise, gp160, which is normally modified by a preponderance of high-mannose/hybrid glycans, could acquire more complex, Endo Hf-resistant carbohydrates after BFA induces the relocation of glycosylation enzymes that are otherwise localized in later Golgi compartments. Similar effects of BFA treatment on processing and glycosylation have been observed for the envelope glycoproteins of vesicular stomatitis virus and murine leukemia virus (126,129). Finally, the disassembly of the Golgi cisternae following BFA treatment apparently disrupts the normal connection between the Golgi and virus assembly sites (42). Thus, depending on the specific conditions, BFA treatment leads to a decrease in virion Env levels.

The consequences of co-expression of sCD4-Ig and full-length, membrane-anchored CD4 on Env cleavage and incorporation into virions were similar, indicating that these effects can be mediated by the two N-terminal CD4 domains and do not require CD4 to be membrane-associated. The transport of intracellular Env-CD4 complexes via the Golgi bypass pathway (42) potentially leads to the exposure of CD4-induced conformations of Env, which are vulnerable to antibody-dependent cellular cytotoxicity (ADCC), on the infected cell surface (88,89). The removal of these Env-CD4 complexes from the infected cell surface is promoted by the HIV-1 Vpu and Nef proteins (88,136–139).

The insights gained from these studies may guide the development of approaches that interrupt Env transport through the secretory pathway and/or interfere with Env cleavage or glycosylation.

## MATERIALS & METHODS

### Cell lines

HEK293T and TZM-bl cells (ATCC) were cultured in Dulbecco modified Eagle medium (DMEM) supplemented with 10% fetal bovine serum (FBS) and 100 µg/ml penicillin/streptomycin (Life Technologies). TZM-bl cells were supplied by John Kappes and Xiaoyun Wu (University of Alabama at Birmingham) through the NIH HIV Reagent Program (140).

### Antibodies (Abs) and CD4 variants

Poorly neutralizing antibodies (pNAbs) used in this study include A32, 17b, b6, 19b and 902090. Broadly neutralizing antibodies (bNAbs) used in this study include 3BNC117, PGDM1400, PGT145, PGT121, PGT151, 35O22 and 10E8. Ab-expressing plasmids were obtained from Dennis Burton (Scripps), Peter Kwong and John Mascola (Vaccine Research Center), James Robinson (Tulane University), Barton Haynes (Duke University) and Hermann Katinger (Polymun) and the NIH HIV Reagent Program. Purified Abs were produced by transfection of Expi293 cells (Thermo Fisher Scientific) with paired heavy-and light-chain Ab-expressing gWiz vectors (aldevron) at a 1:1 ratio. Abs were purified from the transfected cell supernatants with rProtein A-Sepharose (GE Healthcare).

Soluble CD4-Ig (sCD4-Ig) encodes the two N-terminal domains of human CD4 fused to an Ab Fc (141–143). Full-length human CD4 was expressed from a previously described plasmid (144).

### HIV-1 Envs

The HIV-1_AD8_ and HIV-1_AE.2_ Envs were expressed by pNL4-3 infectious proviral clones (92,130,145,146). Plasmids containing the infectious molecular proviral clones of HIV-1_YU2_ (clade B) (120), HIV-1_CH058_ (clade B transmitted/founder) (121) and HIV-1_MJ4_ (clade C) (122) were obtained through the NIH HIV Reagent Program.

### Virus infectivity and neutralization

The effects of Abs on HIV-1 infectivity were measured in two assay formats:

1) Co-expression of HIV-1 and Abs in the same cell – HEK293T cells were cotransfected with 150 ng pNL4-3-AD8 (or with infectious molecular proviral clones encoding Envs from other HIV-1 strains) and 150 ng each of gWiz vectors (aldevron) expressing the Ab heavy chain and light chain, using 0.75 microliters Lipofectamine 3000 Transfection Reagent (Invitrogen). Six to eight hours after transfection, the cell culture medium was replaced with fresh medium. Forty-eight hours after transfection, the infectious virus titer in the HEK293T cell medium was determined on TZM-bl target cells (140,146). Forty-eight hours after incubation with the virus-containing cell supernatants, the TZM-bl cells were lysed and luciferase activity measured.
2) Addition of purified Abs to the medium of virus-producing cells – HEK293T cells were transfected with 150 ng plasmid containing infectious HIV-1 proviral clones. Six to eight hours after transfection, the cell culture medium was replaced with fresh medium containing purified Abs at a concentration of 10 µg/ml. Approximately 40 hours later, the cell supernatants were collected and clarified by low-speed centrifugation (1500 x g at room temperature for 10 minutes). Different amounts of clarified cell supernatants were added to TZM-bl cells. Forty-eight hours later, the TZM-bl cells were lysed and luciferase activity measured.

### Characterization of Env in virus-producing cells and virions

Cells expressing HIV-1 and virions in the supernatants of these cultures were derived by transfection of HEK293T cells, as described above. Six to eight hours after transfection, the medium was changed. In some experiments, the fresh medium contained brefeldin A (BFA) at various concentrations. At 48 hours after transfection, the HEK293T cells were harvested and lysed in 1.5% Cymal-5. Cell supernatants were clarified at 1500 x g for 10 minutes at room temperature. Virus particles were then pelleted at 14,000 x g for 1 hour at 4°C and subsequently lysed in 1.5% Cymal-5. In some cases, cell and virus particle lysates were treated with PNGase F and Endo Hf (New England Biolabs) following the manufacturer’s protocol for 1 hour at 37°C prior to Western blotting.

### Western blotting

Cell and virion lysates were boiled and analyzed by Western blotting using a nitrocellulose membrane and wet transfer (350 mA, 75 min, Bio-Rad). Western blots were developed with 1:2500 goat anti-gp120 polyclonal Ab (Invitrogen), 1:2500 4E10 anti-gp41 Ab (Polymun), 1:5000 rabbit anti-HIV-1 Gag p55/p24/p17 Ab (Abcam), mouse anti-gapdh Ab (Millipore), and/or mouse anti-actin Ab (Invitrogen). The respective HRP-conjugated secondary Abs were 1:2500 rabbit anti-goat Ab (Invitrogen), 1:2500 goat anti-human Ab (Invitrogen), 1:5000 goat anti-rabbit Ab (Sigma-Aldrich), 1:10000 goat anti-mouse Ab (Jackson ImmunoResearch), and 1:10000 goat anti-mouse Ab (Jackson ImmunoResearch).

### Biotinylation of cell-surface Env

Proteins on the surface of Env-expressing HEK293T cells were biotinylated with the Pierce Cell Surface Protein Isolation Kit (Thermo Fisher Scientific). Briefly, the cells were incubated with 2 mM sulfo-NHS-SS-biotin at room temperature for 15 minutes. The cells were then washed twice with 1x glycine in PBS and lysed. The biotinylated proteins in the lysates were affinity-purified on NeutrAvidin-Agarose beads at 4°C for 1 hour and then Western blotted, as described above.

### Immunoprecipitation of cell-surface Env and Env:CD4 complexes

293T cells transiently expressing HIV-1_AD8_ Env, full-length human CD4 or both proteins were analyzed for cell-surface expression of Env and Env:CD4 complexes. Cells were washed twice with 1x PBS/5% FBS. The cells were then incubated with 5 µg/ml OKT4 anti-CD4 antibody (Thermo Fisher Scientific) or a mixture of the 2G12 and PGT121 antibodies against gp120 for 1 hour at 4°C. After four washes in 1x PBS/5% FBS, the cells were lysed in NP-40 lysis buffer (1% NP-40, 0.5 M NaCl, 10 mM Tris [pH 7.5]) for 5 minutes on ice. The lysates were cleared by centrifugation at 13,200 x g for 10 min at 4°C, and the clarified supernatants were incubated with protein A-Sepharose beads for 1 h at room temperature. The beads were pelleted (1,000 rpm x 1 min) and washed three times with wash buffer [20 mM Tris-HCl (pH 8.0), 100 mM (NH_4_)_2_SO_4_, 1 M NaCl, and 0.5% NP-40]. The beads were suspended in 2x lithium dodecyl sulfate (LDS) sample buffer, boiled, and analyzed by Western blotting with 1:2,000 goat anti-gp120 polyclonal antibody (Thermo Fisher Scientific) and 1:2,000 HRP-conjugated rabbit anti-goat IgG (Thermo Fisher Scientific). For analysis of total Env expression in the cells, clarified cell lysates were prepared from cells that were not incubated with antibodies.

The clarified cell lysates were analyzed by Western blotting for Env and actin as described above and serve as the input samples.

## ACKNOWLEDGMENTS

We thank Ms. Elizabeth Carpelan for manuscript preparation. Antibodies against HIV-1 were kindly supplied by Dennis Burton (Scripps), Peter Kwong and John Mascola (Vaccine Research Center NIH), Barton Haynes (Duke University), Hermann Katinger (Polymun), James Robinson (Tulane University), and Marshall Posner (Mount Sinai Medical Center). We thank the NIH HIV Reagent Program for providing reagents.

This work was supported by grants from the National Institutes of Health (grants AI145547, AI124982, AI129017, AI164562, AI150471, AI148379, AI150322, AI129769 and AI176904), by an HIV Cure Research Grant from Gilead Sciences, and by a gift from the late William F. McCarty-Cooper.

## REFERENCES

1. Wyatt R, Sodroski J. 1998. The HIV-1 envelope glycoproteins: fusogens, antigens, and immunogens. Science 280:1884–88.

2. Karlsson Hedestam GB, Fouchier RA, Phogat S, Burton DR, Sodroski J, Wyatt RT. 2008. The challenges of eliciting neutralizing antibodies to HIV-1 and to influenza virus. Nat Rev Microbiol 6:143–55.

3. Hoxie JA. 2010. Toward an antibody-based HIV-1 vaccine. Annu Rev Med 61:135–52.

4. Haynes BF, Shaw GM, Korber B, Kelsoe G, Sodroski J, Hahn BH, Borrow P, McMichael AJ. 2016. HIV-Host interactions: implications for vaccine design. Cell Host Microbe 19:292–303.

5. Fauci AS. 2016. An HIV vaccine: mapping uncharted territory. JAMA 316:143–44.

6. Allan JS, Coligan JE, Barin F, McLane MF, Sodroski JG, Rosen CA, Haseltine WA, Lee TH, Essex M. 1985. Major glycoprotein antigens that induce antibodies in AIDS patients are encoded by HTLV-III. Science 228:1091–94.

7. Robey WG, Safai B, Oroszlan S, Arthur LO, Gonda MA, Gallo RC, Fischinger PJ. 1985. Characterization of envelope and core structural gene products of HTLV-III with sera from AIDS patients. Science 228:593–95.

8. Klatzmann D, Champagne E, Chamaret S, Gruest J, Guetard D, Hercend T, Gluckman JC, Montagnier L. 1984. T-lymphocyte T4 molecule behaves as the receptor for human retrovirus LAV. Nature 312:767–68.

9. Dalgleish AG, Beverley PC, Clapham PR, Crawford DH, Greaves MF, Weiss RA. 1984. The CD4 (T4) antigen is an essential component of the receptor for the AIDS retrovirus. Nature 312:763–67.

10. Wu L, Gerard NP, Wyatt R, Choe H, Parolin C, Ruffing N, Borsetti A, Cardoso AA, Desjardin E, Newman W, Gerard C, Sodroski J. 1996. CD4-induced interaction of primary HIV-1 gp120 glycoproteins with the chemokine receptor CCR-5. Nature 384:179–83.

11. Trkola A, Dragic T, Arthos J, Binley JM, Olson WC, Allaway GP, Cheng-Mayer C, Robinson J, Maddon PJ, Moore JP. 1996. CD4-dependent, antibody-sensitive interactions between HIV-1 and its co-receptor CCR-5. Nature 384:184–87.

12. Choe H, Farzan M, Sun Y, Sullivan N, Rollins B, Ponath PD, Wu L, Mackay CR, LaRosa G, Newman W, Gerard N, Gerard C, Sodroski J. 1996. The beta-chemokine receptors CCR3 and CCR5 facilitate infection by primary HIV-1 isolates. Cell 85:1135–48.

13. Deng H, Liu R, Ellmeier W, Choe S, Unutmaz D, Burkhart M, Di Marzio P, Marmon S, Sutton RE, Hill CM, Davis CB, Peiper SC, Schall TJ, Littman DR, Landau NR. 1996. Identification of a major co-receptor for primary isolates of HIV-1. Nature 381:661–66.

14. Dragic T, Litwin V, Allaway GP, Martin SR, Huang Y, Nagashima KA, Cayanan C, Maddon PJ, Koup RA, Moore JP, Paxton WA. 1996. HIV-1 entry into CD4+ cells is mediated by the chemokine receptor CC-CKR-5. Nature 381:667–73.

15. Doranz BJ, Rucker J, Yi Y, Smyth RJ, Samson M, Peiper SC, Parmentier M, Collman RG, Doms RW. 1996. A dual-tropic primary HIV-1 isolate that uses fusin and the beta-chemokine receptors CKR-5, CKR-3, and CKR-2b as fusion cofactors. Cell 85:1149–58.

16. Feng Y, Broder CC, Kennedy PE, Berger EA. 1996. HIV-1 entry cofactor: functional cDNA cloning of a seven-transmembrane, G protein-coupled receptor. Science 272:872–77.

17. Alkhatib G, Combadiere C, Broder CC, Feng Y, Kennedy PE, Murphy PM, Berger EA. 1996. CC CKR5: a RANTES, MIP-1alpha, MIP-1beta receptor as a fusion cofactor for macrophage-tropic HIV-1. Science 272:1955–58.

18. Chan DC, Fass D, Berger JM, Kim PS. 1997. Core structure of gp41 from the HIV envelope glycoprotein. Cell 89:26373.

19. Weissenhorn W, Dessen A, Harrison SC, Skehel JJ, Wiley DC. 1997. Atomic structure of the ectodomain from HIV-1 gp41. Nature 387:426–30.

20. Lu M, Blacklow SC, Kim PS. 1995. A trimeric structural domain of the HIV-1 transmembrane glycoprotein. Nat Struct Biol 2:1075–82.

21. Dalgleish AG, Beverley PC, Clapham PR, Crawford DH, Greaves MF, Weiss RA. 1984. The CD4 (T4) antigen is an essential component of the receptor for the AIDS retrovirus. Nature 312:763–67.

22. Fennie C, Lasky LA. 1989. Model for intracellular folding of the human immunodeficiency virus type 1 gp120. J Virol 63:639–46.

23. Li Y, Luo L, Thomas DY, Kang CY. 2000. The HIV-1 Env protein signal sequence retards its cleavage and down-regulates the glycoprotein folding. Virology 272:417–28.

24. Willey RL, Bonifacino JS, Potts BJ, Martin MA, Klausner RD. 1988. Biosynthesis, cleavage, and degradation of the human immunodeficiency virus 1 envelope glycoprotein gp160. Proc Natl Acad Sci U S A 85:9580–84.

25. Earl PL, Moss B, Doms RW. 1991. Folding, interaction with GRP78-BiP, assembly, and transport of the human immunodeficiency virus type 1 envelope protein. J Virol 65:2047–55.

26. Bosch V, Pawlita M. 1990. Mutational analysis of the human immunodeficiency virus type 1 env gene product proteolytic cleavage site. J Virol 64:2337–44.

27. Decroly E, Vandenbranden M, Ruysschaert JM, Cogniaux J, Jacob GS, Howard SC, Marshall G, Kompelli A, Basak A, Jean F, Lazuref C, Bedannet S, Chrétien M, Day R, Seidah NG. 1994. The convertases furin and PC1 can both cleave the human immunodeficiency virus (HIV)-1 envelope glycoprotein gp160 into gp120 (HIV-1 SU) and gp41 (HIV-I TM). J Biol Chem 269:12240–47.

28. Fenouillet E, Gluckman JC. 1992. Immunological analysis of human immunodeficiency virus type 1 envelope glycoprotein proteolytic cleavage. Virology 187:825–28.

29. Hallenberger S, Bosch V, Angliker H, Shaw E, Klenk HD, Garten W. 1992. Inhibition of furin-mediated cleavage activation of HIV-1 glycoprotein gp160. Nature 360:358–61.

30. Dewar RL, Natarajan V, Vasudevachari MB, Salzman NP. 1989. Synthesis and processing of human immunodeficiency virus type 1 envelope proteins encoded by a recombinant human adenovirus. J Virol 63:129–36.

31. Dewar RL, Vasudevachari MB, Natarajan V, Salzman NP. 1989. Biosynthesis and processing of human immunodeficiency virus type 1 envelope glycoproteins: effects of monensin on glycosylation and transport. J Virol 63:2452–56.

32. Merkle RK, Helland DE, Welles JL, Shilatifard A, Haseltine WA, Cummings RD. 1991. gp160 of HIV-I synthesized by persistently infected Molt-3 cells is terminally glycosylated: evidence that cleavage of gp160 occurs subsequent to oligosaccharide processing. Arch Biochem Biophys 290:248–57.

33. Kantanen ML, Leinikki P, Kuismanen E. 1995. Endoproteolytic cleavage of HIV-1 gp160 envelope precursor occurs after exit from the trans-Golgi network (TGN). Arch Virol 140:1441–49.

34. Pfeiffer T, Zentgraf H, Freyaldenhoven B, Bosch V. 1997. Transfer of endoplasmic reticulum and Golgi retention signals to human immunodeficiency virus type 1 gp160 inhibits intracellular transport and proteolytic processing of viral glycoprotein but does not influence the cellular site of virus particle budding. J Gen Virol 78:1745–53.

35. Miranda L, Wolf J, Pichuantes S, Duke R, Franzusoff A. 1996. Isolation of the human PC6 gene encoding the putative host protease for HIV-1 gp160 processing in CD4+ T lymphocytes. Proc Natl Acad Sci U S A 93:7695–7700.

36. Ohnishi Y, Shioda T, Nakayama K, Iwata S, Gotoh B, Hamaguchi M, Nagai Y. 1994. A furin-defective cell line is able to process correctly the gp160 of human immunodeficiency virus type 1. J Virol 68:4075–79.

37. Stein BS, Engleman EG. 1990. Intracellular processing of the gp160 HIV-1 envelope precursor. Endoproteolytic cleavage occurs in a cis or medial compartment of the Golgi complex. J Biol Chem 265:2640–49.

38. Helseth E, Olshevsky U, Furman C, Sodroski J. 1991. Human immunodeficiency virus type 1 gp120 envelope glycoprotein regions important for association with the gp41 transmembrane glycoprotein. J Virol 65:2119–23.

39. Gabuzda D, Olshevsky U, Bertani P, Haseltine WA, Sodroski J. 1991. Identification of membrane anchorage domains of the HIV-1 gp160 envelope glycoprotein precursor. J Acquir Immune Defic Syndr 4:34–40.

40. Haffar OK, Dowbenko DJ, Berman PW. 1988. Topogenic analysis of the human immunodeficiency virus type 1 envelope glycoprotein, gp160, in microsomal membranes. J Cell Biol 107:1677–87.

41. Dev J, Park D, Fu Q, Chen J, Ha HJ, Ghantous F, Herrmann T, Chang W, Liu Z, Frey G, Seaman MS, Chen B, Chou JJ. 2016. Structural basis for membrane anchoring of HIV-1 envelope spike. Science 353:172–75.

42. Zhang S, Nguyen HT, Ding H, Wang J, Zou S, Liu L, Guha D, Gabuzda D, Ho DD, Kappes JC, Sodroski J. 2021. Dual pathways of human immunodeficiency virus type 1 envelope glycoprotein trafficking modulate the selective exclusion of uncleaved oligomers from virions. J Virol 95:e01369–20.

43. Munro JB, Gorman J, Ma X, Zhou Z, Arthos J, Burton DR, Koff WC, Courter JR, Smith AB III, Kwong PD, Blanchard SC, Mothes W. 2014. Conformational dynamics of single HIV-1 envelope trimers on the surface of native virions. Science 346:759–63.

44. Herschhorn A, Ma X, Gu C, Ventura JD, Castillo-Menendez L, Melillo B, Terry DS, Smith AB III, Blanchard SC, Munro JB, Mothes W, Finzi A, Sodroski J. 2016. Release of gp120 restraints leads to an entry-competent intermediate state of the HIV-1 envelope glycoproteins. MBio 7:e01598–16.

45. Ma X, Terry DS, Gorman J, Hong X, Zhou Z, Zhao H, Altman RB, Arthos J, Kwong PD, Blanchard SC, Mothes W, Munro JB. 2018. HIV-1 Env trimer opens through an asymmetric intermediate in which individual protomers adopt distinct conformations. eLife 7:e34271.

46. Haim H, Strack B, Kassa A, Madani N, Wang L, Courter JR, Princiotto A, McGee K, Pacheco B, Seaman MS, Smith AB III, Sodroski J. 2011. Contribution of intrinsic reactivity of the HIV-1 envelope glycoproteins to CD4-independent infection and global inhibitor sensitivity. PLoS Pathog 7:e1002101.

47. Furuta RA, Wild CT, Weng Y, Weiss CD. 1998. Capture of an early fusion-active conformation of HIV-1 gp41. Nat Struct Biol 5:276–79.

48. Koshiba T, Chan DC. 2003. The prefusogenic intermediate of HIV-1 gp41 contains exposed C-peptide regions. J Biol Chem 278:7573–79.

49. He Y, Vassell R, Zaitseva M, Nguyen N, Yang Z, Weng Y, Weiss CD. 2003. Peptides trap the human immunodeficiency virus type 1 envelope glycoprotein fusion intermediate at two sites. J Virol 77:1666–71.

50. Si Z, Madani N, Cox JM, Chruma JJ, Klein JC, Schon A, Phan N, Wang L, Biorn AC, Cocklin S, Chaiken I, Freire E, Smith AB III, Sodroski JG. 2004. Small-molecule inhibitors of HIV-1 entry block receptor-induced conformational changes in the viral envelope glycoproteins. Proc Natl Acad Sci U S A 101:5036–41.

51. Herschhorn A, Gu C, Moraca F, Ma X, Farrell M, Smith AB III, Pancera M, Kwong PD, Schon A, Freire E, Abrams C, Blanchard SC, Mothes W, Sodroski JG. 2017. The beta20-beta21 of gp120 is a regulatory switch for HIV-1 Env conformational transitions. Nat Commun 8:1049.

52. Castillo-Menendez LR, Nguyen HT, Sodroski J. 2019. Conformational differences between functional human immunodeficiency virus (HIV-1) envelope glycoprotein trimers and stabilized soluble trimers. J Virol 93:e01709–18.

53. Ivan B, Sun Z, Subbaraman H, Friedrich N, Trkola A. 2019. CD4 occupancy triggers sequential pre-fusion conformational states of the HIV-1 envelope trimer with relevance for broadly neutralizing antibody activity. PLoS Biol 17:e3000114.

54. Kuhmann SE, Platt EJ, Kozak SL, Kabat D. 2000. Cooperation of multiple CCR5 coreceptors is required for infections by human immunodeficiency virus type 1. J Virol 74:7005–15.

55. Melikyan GB, Markosyan RM, Hemmati H, Delmedico MK, Lambert DM, Cohen FS. 2000. Evidence that the transition of HIV-1 gp41 into a six-helix bundle, not the bundle configuration, induces membrane fusion. J Cell Biol 151:413–23.

56. Wilen CB, Tilton JC, Doms RW. 2012. Molecular mechanisms of HIV entry. In: Rossmann M, Rao V (eds.), Viral Molecular Machines. Advances in Experimental Medicine and Biology, vol 726. Springer, Boston, MA.

57. Fouts TR, Binley JM, Trkola A, Robinson JE, Moore JP. 1997. Neutralization of the human immunodeficiency virus type 1 primary isolate JR-FL by human monoclonal antibodies correlates with antibody binding to the oligomeric form of the envelope glycoprotein complex. J Virol 71:2779–85.

58. York J, Follis KE, Trahey M, Nyambi PN, Zolla-Pazner S, Nunberg JH. 2001. Antibody binding and neutralization of primary and T-cell line-adapted isolates of human immunodeficiency virus type 1. J Virol 75:2741–52.

59. Ren X, Sodroski J, Yang X. 2005. An unrelated monoclonal antibody neutralizes human immunodeficiency virus type 1 by binding to an artificial epitope engineered in a functionally neutral region of the viral envelope glycoproteins. J Virol 79:5616–24.

60. Yang X, Lipchina I, Cocklin S, Chaiken I, Sodroski J. 2006. Antibody binding is a dominant determinant of the efficiency of human immunodeficiency virus type 1 neutralization. J Virol 80:11404–408.

61. Haim H, Salas I, McGee K, Eichelberger N, Winter E, Pacheco B, Sodroski J. 2013. Modeling virus-and antibody-specific factors to predict human immunodeficiency virus neutralization efficiency. Cell Host Microbe 14:547–58.

62. Guttman M, Cupo A, Julien JP, Sanders RW, Wilson IA, Moore JP, Lee KK. 2015. Antibody potency relates to the ability to recognize the closed, pre-fusion form of HIV Env. Nat Commun 6:6144.

63. Kwong PD, Wyatt R, Robinson J, Sweet RW, Sodroski J, Hendrickson WA. 1998. Structure of an HIV gp120 envelope glycoprotein in complex with the CD4 receptor and a neutralizing human antibody. Nature 393:648–59.

64. Wyatt R, Kwong PD, Desjardins E, Sweet RW, Robinson J, Hendrickson WA, Sodroski JG. 1998. The antigenic structure of the HIV gp120 envelope glycoprotein. Nature 393:705–11.

65. Kwong PD, Doyle ML, Casper DJ, Cicala C, Leavitt SA, Majeed S, Steenbeke TD, Venturi M, Chaiken I, Fung M, Katinger H, Parren PW, Robinson J, Van Ryk D, Wang L, Burton DR, Freire E, Wyatt R, Sodroski J, Hendrickson WA, Arthos J. 2002. HIV-1 evades antibody-mediated neutralization through conformational masking of receptor-binding sites. Nature 420:678–82.

66. Kuiken C, Foley B, Marx P, Wolinsky S, Leitner T, Hahn B, McCutchan F, Korber B. HIV Sequence Compendium 2013. Los Alamos HIV Sequence Database.

67. Wei X, Decker JM, Wang S, Hui H, Kappes JC, Wu X, Salazar-Gonzalez JF, Salazar MG, Kilby JM, Saag MS, Komarova NL, Nowak MA, Hahn BH, Kwong PD, Shaw GM. 2003. Antibody neutralization and escape by HIV-1. Nature 422:307–12.

68. Burton DR, Desrosiers RC, Doms RW, Koff WC, Kwong PD, Moore JP, Nabel GJ, Sodroski J, Wilson IA, Wyatt RT. 2004. HIV vaccine design and the neutralizing antibody problem. Nat Immunol 5:233–36.

69. Moore PL, Ranchobe N, Lambson BE, Gray ES, Cave E, Abrahams MR, Bandawe G, Mlisana K, Abdool Karim SS, Williamson C, Morris L, CAPRISA 002 study, NIAID Center for HIV/AIDS Vaccine Immunology (CHAVI). 2009. Limited neutralizing antibody specificities drive neutralization escape in early HIV-1 subtype C infection. PLoS Pathog 5:e1000598.

70. Wibmer CK, Bhiman JN, Gray ES, Tumba N, Abdool Karim SS, Williamson C, Morris L, Moore PL. 2013. Viral escape from HIV-1 neutralizing antibodies drives increased plasma neutralization breadth through sequential recognition of multiple epitopes and immunotypes. PLoS Pathog 9:e1003738.

71. Gray ES, Taylor N, Wycuff D, Moore PL, Tomaras GD, Wibmer CK, Puren A, DeCamp A, Gilbert PB, Wood B, Montefiori DC, Binley JM, Shaw GM, Haynes BF, Mascola JR, Morris L. 2009. Antibody specificities associated with neutralization breadth in plasma from human immunodeficiency virus type 1 subtype C-infected blood donors. J Virol 83:8925–37.

72. Sather DN, Armann J, Ching LK, Mavrantoni A, Sellhorn G, Caldwell Z, Yu X, Wood B, Self S, Kalams S, Stamatatos L. 2009. Factors associated with the development of cross-reactive neutralizing antibodies during human immunodeficiency virus type 1 infection. J Virol 83:757–69.

73. Klein F, Diskin R, Scheid JF, Gaebler C, Mouquet H, Georgiev IS, Pancera M, Zhou T, Incesu RB, Fu BZ, Gnanapragasam PN, Oliveira TY, Seaman MS, Kwong PD, Bjorkman PJ, Nussenzweig MC. 2013. Somatic mutations of the immunoglobulin framework are generally required for broad and potent HIV-1 neutralization. Cell 153:126–38.

74. Walker LM, Simek MD, Priddy F, Gach JS, Wagner D, Zwick MB, Phogat SK, Poignard P, Burton DR. 2010. A limited number of antibody specificities mediate broad and potent serum neutralization in selected HIV-1 infected individuals. PLoS Pathog 6:e1001028.

75. Gray ES, Madiga MC, Hermanus T, Moore PL, Wibmer CK, Tumba NL, Werner L, Mlisana K, Sibeko S, Williamson C, Abdool Karim SS, Morris L, Team CS. 2011. The neutralization breadth of HIV-1 develops incrementally over four years and is associated with CD4+ T cell decline and high viral load during acute infection. J Virol 85:4828–40.

76. Corti D, Langedijk JP, Hinz A, Seaman MS, Vanzetta F, Fernandez-Rodriguez BM, Silacci C, Pinna D, Jarrossay D, Balla-Jhagjhoorsingh S, Willems B, Zekveld MJ, Dreja H, O’Sullivan E, Pade C, Orkin C, Jeffs SA, Montefiori DC, Davis D, Weissenhorn W, McKnight A, Heeney JL, Sallusto F, Sattentau QJ, Weiss RA, Lanzavecchia A. 2010. Analysis of memory B cell responses and isolation of novel monoclonal antibodies with neutralizing breadth from HIV-1-infected individuals. PLoS One 5:e8805.

77. Wu X, Zhou T, Zhu J, Zhang B, Georgiev I, Wang C, Chen X, Longo NS, Louder M, McKee K, O’Dell S, Perfetto S, Schmidt SD, Shi W, Wu L, Yang Y, Yang ZY, Yang Z, Zhang Z, Bonsignori M, Crump JA, Kapiga SH, Sam NE, Haynes BF, Simek M, Burton DR, Koff WC, Doria-Rose NA, Connors M, Program NCS, Mullikin JC, Nabel GJ, Roederer M, Shapiro L, Kwong PD, Mascola JR. 2011. Focused evolution of HIV-1 neutralizing antibodies revealed by structures and deep sequencing. Science 333:1593–1602.

78. Hraber P, Seaman MS, Bailer RT, Mascola JR, Montefiori DC, Korber BT. 2014. Prevalence of broadly neutralizing antibody responses during chronic HIV-1 infection. AIDS 28:163–69.

79. Castillo-Menendez LR, Witt K, Espy N, Princiotto A, Madani N, Pacheco B, Finzi A, Sodroski J. 2018. Comparison of uncleaved and mature human immunodeficiency virus membrane envelope glycoprotein trimers. J Virol 92:e00277–18.

80. Wang Q, Finzi A, Sodroski J. 2020. The conformational states of the HIV-1 envelope glycoproteins. 2020. Trends Microbiol 28:655–67.

81. Lu M, Ma X, Reichard N, Terry DS, Arthos J, Smith AB III, Sodroski JG, Blanchard SC, Mothes W. 2020. Shedding-resistant HIV-1 envelope glycoproteins adopt downstream conformations that remain responsive to conformation-preferring ligands. J Virol 94:e00597–20.

82. Pancera M, Wyatt R. 2005. Selective recognition of oligomeric HIV-1 primary isolate envelope glycoproteins by potently neutralizing ligands requires efficient precursor cleavage. Virology 332:145–56.

83. Chakrabarti BK, Pancera M, Phogat S, O’Dell S, McKee K, Guenaga J, Robinson J, Mascola J, Wyatt RT. 2011. HIV type 1 Env precursor cleavage state affects recognition by both neutralizing and nonneutralizing gp41 antibodies. AIDS Res Hum Retroviruses 27:877–87.

84. Zou S, Zhang S, Gaffney A, Ding H, Lu M, Grover JR, Farrell M, Nguyen HT, Zhao C, Anang S, Zhao M, Mohammadi M, Blanchard SC, Abrams C, Madani N, Mothes W, Kappes JC, Smith AB III, Sodroski J. 2020. Long-acting BMS-378806 analogues stabilize the State-1 conformation of the human immunodeficiency virus type 1 envelope glycoproteins. J Virol 94:e00148–20.

85. Haim H, Salas I, Sodroski J. 2013. Proteolytic processing of the human immunodeficiency virus envelope glycoprotein precursor decreases conformational flexibility. J Virol 87:1884–89.

86. Zhang S, Wang K, Wang WL, Nguyen HT, Chen S, Lu M, Go EP, Ding H, Steinbock RT, Desaire H, Kappes JC, Sodroski J, Mao Y. 2021. Asymmetric structures and conformational plasticity of the uncleaved full-length human immunodeficiency virus envelope glycoprotein trimer. J Virol 95:e0052921.

87. Hoxie JA, Alpers JD, Rackowski JL, Huebner K, Haggarty BS, Cedarbaum AJ, Reed JC. 1986. Alterations in T4 (CD4) protein and mRNA synthesis in cells infected with HIV. Science 234:1123–27.

88. Veillette M, Desormeaux A, Medjahed H, Gharsallah NE, Coutu M, Baalwa J, Guan Y, Lewis G, Ferrari G, Hahn BH, Haynes BF, Robinson JE, Kaufmann DE, Bonsignori M, Sodroski J, Finzi A. 2014. Interaction with cellular CD4 exposes HIV-1 envelope epitopes targeted by antibody-dependent cell-mediated cytotoxicity. J Virol 88:2633–44.

89. Veillette M, Coutu M, Richard J, Batraville LA, Dagher O, Bernard N, Tremblay C, Kaufmann DE, Roger M, Finzi A. 2015. The HIV-1 gp120 CD4-bound conformation is preferentially targeted by antibody-dependent cellular cytotoxicity-mediating antibodies in sera from HIV-1-infected individuals. J Virol 89:545–51.

90. Witt KC, Castillo-Menendez L, Ding H, Espy N, Zhang S, Kappes JC, Sodroski J. 2017. Antigenic characterization of the human immunodeficiency virus (HIV-1) envelope glycoprotein precursor incorporated into nanodiscs. PLoS One 12:e0170672.

91. Zhou R, Zhang S, Nguyen HT, Ding H, Gaffney A, Kappes JC, Smith AB III, Sodroski JG. 2023. Conformations of human immunodeficiency virus envelope glycoproteins in detergents and styrene-maleic acid lipid particles. J Virol 97:e0032723.

92. Nguyen HT, Wang Q, Anang S, Sodroski JG. 2023. Characterization of the human immunodeficiency virus (HIV-1) envelope glycoprotein conformational states on infectious virus particles. J Virol 97:e0185722.

93. Moore JP, Trkola A, Korber B, Boots LJ, Kessler JA, 2nd, McCutchan FE, Mascola J, Ho DD, Robinson J, Conley AJ. 1995. A human monoclonal antibody to a complex epitope in the V3 region of gp120 of human immunodeficiency virus type 1 has broad reactivity within and outside clade B. J Virol 69:122–30.

94. Thali M, Moore JP, Furman C, Charles M, Ho DD, Robinson J, Sodroski J. 1993. Characterization of conserved human immunodeficiency virus type 1 gp120 neutralization epitopes exposed upon gp120-CD4 binding. J Virol 67:3978–88.

95. Richard J, Pacheco B, Gohain N, Veillette M, Ding S, Alsahafi N, Tolbert WD, Prevost J, Chapleau JP, Coutu M, Jia M, Brassard N, Park J, Courter JR, Melillo B, Martin L, Tremblay C, Hahn BH, Kaufmann DE, Wu X, Smith AB III, Sodroski J, Pazgier M, Finzi A. 2016. Co-receptor binding site antibodies enable CD4-mimetics to expose conserved anti-cluster A ADCC epitopes on HIV-1 envelope glycoproteins. EBioMedicine 12:208–18.

96. Chen L, Kwon YD, Zhou T, Wu X, O’Dell S, Cavacini L, Hessell AJ, Pancera M, Tang M, Xu L, Yang ZY, Zhang MY, Arthos J, Burton DR, Dimitrov DS, Nabel GJ, Posner MR, Sodroski J, Wyatt R, Mascola JR, Kwong PD. 2009. Structural basis of immune evasion at the site of CD4 attachment on HIV-1 gp120. Science 326:1123–27.

97. McCoy LE, Burton DR. 2017. Identification and specificity of broadly neutralizing antibodies against HIV. Immunol Rev 275:11–20.

98. Sok D, Burton DR. 2018. Recent progress in broadly neutralizing antibodies to HIV. Nat Immunol 19:1179–88.

99. Haynes BF, Burton DR, Mascola JR. 2019. Multiple roles for HIV broadly neutralizing antibodies. Sci Transl Med 11:eaaz2686.

100. Ward AB, Wilson IA. 2017. The HIV-1 envelope glycoprotein structure: nailing down a moving target. Immunol Rev 275:21–32.

101. Kwong PD, Mascola JR. 2018. HIV-1 vaccines based on antibody identification, B cell ontogeny, and epitope structure. Immunity 48:855–71.

102. Zhou T, Georgiev I, Wu X, Yang ZY, Dai K, Finzi A, Kwon YD, Scheid JF, Shi W, Xu L, Yang Y, Zhu J, Nussenzweig MC, Sodroski J, Shapiro L, Nabel GJ, Mascola JR, Kwong PD. 2010. Structural basis for broad and potent neutralization of HIV-1 by antibody VRC01. Science 329:811–17.

103. Zhou T, Lynch RM, Chen L, Acharya P, Wu X, Doria-Rose NA, Joyce MG, Lingwood D, Soto C, Bailer RT, Ernandes MJ, Kong R, Longo NS, Louder MK, McKee K, O’Dell S, Schmidt SD, Tran L, Yang Z, Druz A, Luongo TS, Moquin S, Srivatsan S, Yang Y, Zhang B, Zheng A, Pancera M, Kirys T, Georgiev IS, Gindin T, Peng HP, Yang AS, Program NCS, Mullikin JC, Gray MD, Stamatatos L, Burton DR, Koff WC, Cohen MS, Haynes BF, Casazza JP, Connors M, Corti D, Lanzavecchia A, Sattentau QJ, Weiss RA, West AP, Jr., Bjorkman PJ, Scheid JF, Nussenzweig MC, et al. 2015. Structural repertoire of HIV-1-neutralizing antibodies targeting the CD4 supersite in 14 donors. Cell 161:1280–92.

104. Walker LM, Phogat SK, Chan-Hui PY, Wagner D, Phung P, Goss JL, Wrin T, Simek MD, Fling S, Mitcham JL, Lehrman JK, Priddy FH, Olsen OA, Frey SM, Hammond PW, Protocol GPI, Kaminsky S, Zamb T, Moyle M, Koff WC, Poignard P, Burton DR. 2009. Broad and potent neutralizing antibodies from an African donor reveal a new HIV-1 vaccine target. Science 326:285–89.

105. Pancera M, Shahzad-Ul-Hussan S, Doria-Rose NA, McLellan JS, Bailer RT, Dai K, Loesgen S, Louder MK, Staupe RP, Yang Y, Zhang B, Parks R, Eudailey J, Lloyd KE, Blinn J, Alam SM, Haynes BF, Amin MN, Wang LX, Burton DR, Koff WC, Nabel GJ, Mascola JR, Bewley CA, Kwong PD. 2013. Structural basis for diverse N-glycan recognition by HIV-1-neutralizing V1-V2-directed antibody PG16. Nat Struct Mol Biol 20:804–13.

106. Walker LM, Huber M, Doores KJ, Falkowska E, Pejchal R, Julien JP, Wang SK, Ramos A, Chan-Hui PY, Moyle M, Mitcham JL, Hammond PW, Olsen OA, Phung P, Fling S, Wong CH, Phogat S, Wrin T, Simek MD, Protocol GPI, Koff WC, Wilson IA, Burton DR, Poignard P. 2011. Broad neutralization coverage of HIV by multiple highly potent antibodies. Nature 477:466–70.

107. Blattner C, Lee JH, Sliepen K, Derking R, Falkowska E, de la Pena AT, Cupo A, Julien JP, van Gils M, Lee PS, Peng W, Paulson JC, Poignard P, Burton DR, Moore JP, Sanders RW, Wilson IA, Ward AB. 2014. Structural delineation of a quaternary, cleavage-dependent epitope at the gp41-gp120 interface on intact HIV-1 Env trimers. Immunity 40:669–80.

108. Huang J, Kang BH, Pancera M, Lee JH, Tong T, Feng Y, Imamichi H, Georgiev IS, Chuang GY, Druz A, Doria-Rose NA, Laub L, Sliepen K, van Gils MJ, de la Pena AT, Derking R, Klasse PJ, Migueles SA, Bailer RT, Alam M, Pugach P, Haynes BF, Wyatt RT, Sanders RW, Binley JM, Ward AB, Mascola JR, Kwong PD, Connors M. 2014. Broad and potent HIV-1 neutralization by a human antibody that binds the gp41-gp120 interface. Nature 515:138–42.

109. Sun ZY, Oh KJ, Kim M, Yu J, Brusic V, Song L, Qiao Z, Wang JH, Wagner G, Reinherz EL. 2008. HIV-1 broadly neutralizing antibody extracts its epitope from a kinked gp41 ectodomain region on the viral membrane. Immunity 28:52–63.

110. Huang J, Ofek G, Laub L, Louder MK, Doria-Rose NA, Longo NS, Imamichi H, Bailer RT, Chakrabarti B, Sharma SK, Alam SM, Wang T, Yang Y, Zhang B, Migueles SA, Wyatt R, Haynes BF, Kwong PD, Mascola JR, Connors M. 2012. Broad and potent neutralization of HIV-1 by a gp41-specific human antibody. Nature 491:406–12.

111. Irimia A, Serra AM, Sarkar A, Jacak R, Kalyuzhniy O, Sok D, Saye-Francisco KL, Schiffner T, Tingle R, Kubitz M, Adachi Y, Stanfield RL, Deller MC, Burton DR, Schief WR, Wilson IA. 2017. Lipid interactions and angle of approach to the HIV-1 viral membrane of broadly neutralizing antibody 10E8: Insights for vaccine and therapeutic design. PLoS Pathog 13:e1006212.

112. Caskey M, Klein F, Lorenzi JC, Seaman MS, West AP, Jr., Buckley N, Kremer G, Nogueira L, Braunschweig M, Scheid JF, Horwitz JA, Shimeliovich I, Ben-Avraham S, Witmer-Pack M, Platten M, Lehmann C, Burke LA, Hawthorne T, Gorelick RJ, Walker BD, Keler T, Gulick RM, Fatkenheuer G, Schlesinger SJ, Nussenzweig MC. 2015. Viraemia suppressed in HIV-1-infected humans by broadly neutralizing antibody 3BNC117. Nature 522:487–91.

113. Sok D, van Gils MJ, Pauthner M, Julien JP, Saye-Francisco KL, Hsueh J, Briney B, Lee JH, Le KM, Lee PS, Hua Y, Seaman MS, Moore JP, Ward AB, Wilson IA, Sanders RW, Burton DR. 2014. Recombinant HIV envelope trimer selects for quaternary-dependent antibodies targeting the trimer apex. Proc Natl Acad Sci U S A 111:17624–29.

114. Roben P, Moore JP, Thali M, Sodroski J, Barbas CF, 3rd, Burton DR. 1994. Recognition properties of a panel of human recombinant Fab fragments to the CD4 binding site of gp120 that show differing abilities to neutralize human immunodeficiency virus type 1. J Virol 68:4821–28.

115. Madani N, Princiotto AM, Easterhoff D, Bradley T, Luo K, Williams WB, Liao HX, Moody MA, Phad GE, Vazquez Bernat N, Melillo B, Santra S, Smith AB III, Karlsson Hedestam GB, Haynes B, Sodroski J. 2016. Antibodies elicited by multiple envelope glycoprotein immunogens in primates neutralize primary human immunodeficiency viruses (HIV-1) sensitized by CD4-mimetic compounds. J Virol 90:5031–46.

116. Murin CD, Fusco ML, Bornholdt ZA, Qiu X, Olinger GG, Zeitlin L, Kobinger GP, Ward AB, Saphire EO. 2014. Structures of protective antibodies reveal sites of vulnerability on Ebola virus. Proc Natl Acad Sci U S A 111:17182–87.

117. Audet J, Wong G, Wang H, Lu G, Gao GF, Kobinger G, Qiu X. 2014. Molecular characterization of the monoclonal antibodies composing ZMAb: a protective cocktail against Ebola virus. Sci Rep 4:6881.

118. Houser KV, Gretebeck L, Ying T, Wang Y, Vogel L, Lamirande EW, Bock KW, Moore IN, Dimitrov DS, Subbarao K. 2016. Prophylaxis with a Middle East respiratory syndrome coronavirus (MERS-CoV)-specific human monoclonal antibody protects rabbits from MERS-CoV infection. J Infect Dis 213:1557–61.

119. Yuan M, Wu NC, Zhu X, Lee CD, So RTY, Lv H, Mok CKP, Wilson IA. 2020. A highly conserved cryptic epitope in the receptor binding domains of SARS-CoV-2 and SARS-CoV. Science 368:630–33.

120. Li Y, Kappes JC, Conway JA, Price RW, Shaw GM, Hahn BH. 1991. Molecular characterization of human immunodeficiency virus type 1 cloned directly from uncultured human brain tissue: identification of replication-competent and -defective viral genomes. J Virol 65:3973–85.

121. Liao HX, Lynch R, Zhou T, Gao F, Alam SM, Boyd SD, Fire AZ, Roskin KM, Schramm CA, Zhang Z, Zhu J, Shapiro L, Program NCS, Mullikin JC, Gnanakaran S, Hraber P, Wiehe K, Kelsoe G, Yang G, Xia SM, Montefiori DC, Parks R, Lloyd KE, Scearce RM, Soderberg KA, Cohen M, Kamanga G, Louder MK, Tran LM, Chen Y, Cai F, Chen S, Moquin S, Du X, Joyce MG, Srivatsan S, Zhang B, Zheng A, Shaw GM, Hahn BH, Kepler TB, Korber BT, Kwong PD, Mascola JR, Haynes BF. 2013. Co-evolution of a broadly neutralizing HIV-1 antibody and founder virus. Nature 496:469–76.

122. Ndung’u T, Renjifo B, Essex M. 2001. Construction and analysis of an infectious human Immunodeficiency virus type 1 subtype C molecular clone. J Virol 75:4964–72.

123. Moore JP, McKeating JA, Weiss RA, Sattentau QJ. 1990. Dissociation of gp120 from HIV-1 virions induced by soluble CD4. Science 250:1139–42.

124. Fujiwara T, Oda K, Yokota S, Takatsuki A, Ikehara Y. 1988. Brefeldin A causes disassembly of the Golgi complex and accumulation of secretory proteins in the endoplasmic reticulum. J Biol Chem 263:18545–52.

125. Ulmer JB, Palade GE. 1989. Targeting and processing of glycophorins in murine erythroleukemia cells: use of brefeldin A as a perturbant of intracellular traffic. Proc Natl Acad Sci U S A 86:6992–96.

126. Doms RW, Russ G, Yewdell JW. 1989. Brefeldin A redistributes resident and itinerant Golgi proteins to the endoplasmic reticulum. J Cell Biol 109:61–72.

127. Lippincott-Schwartz J, Yuan LC, Bonifacino JS, Klausner RD. 1989. Rapid redistribution of Golgi proteins into the ER in cells treated with brefeldin A: evidence for membrane cycling from Golgi to ER. Cell 56:801–13.

128. Lippincott-Schwartz J, Donaldson JG, Schweizer A, Berger EG, Hauri HP, Yuan LC, Klausner RD. 1990. Microtubule-dependent retrograde transport of proteins into the ER in the presence of brefeldin A suggests an ER recycling pathway. Cell 60:821–36.

129. Ulmer JB, Palade GE. 1991. Effects of brefeldin A on the processing of viral envelope glycoproteins in murine erythroleukemia cells. J Biol Chem 266:9173–79.

130. Nguyen HT, Qualizza A, Anang S, Zhao M, Zou S, Zhou R, Wang Q, Zhang S, Deshpande A, Ding H, Chiu TJ, Smith AB III, Kappes JC, Sodroski JG. 2022. Functional and highly cross-linkable HIV-1 envelope glycoproteins enriched in a pretriggered conformation. J Virol 96:e0166821.

131. Ruprecht CR, Krarup A, Reynell L, Mann AM, Brandenberg OF, Berlinger L, Abela IA, Regoes RR, Gunthard HF, Rusert P, Trkola A. 2011. MPER-specific antibodies induce gp120 shedding and irreversibly neutralize HIV-1. J Exp Med 208:439–54.

132. Li Y, O’Dell S, Walker LM, Wu X, Guenaga J, Feng Y, Schmidt SD, McKee K, Louder MK, Ledgerwood JE, Graham BS, Haynes BF, Burton DR, Wyatt RT, Mascola JR. 2011. Mechanism of neutralization by the broadly neutralizing HIV-1 monoclonal antibody VRC01. J Virol 85:8954–67.

133. Haim H, Si Z, Madani N, Wang L, Courter JR, Princiotto A, Kassa A, DeGrace M, McGee-Estrada K, Mefford M, Gabuzda D, Smith AB III, Sodroski J. 2009. Soluble CD4 and CD4-mimetic compounds inhibit HIV-1 infection by induction of a short-lived activated state. PLoS Pathog 5:e1000360.

134. Yang X, Kurteva S, Ren X, Lee S, Sodroski J. 2005. Stoichiometry of envelope glycoprotein trimers in the entry of human immunodeficiency virus type 1. J Virol 79:12132–47.

135. Misumi Y, Misumi Y, Miki K, Takatsuki A, Tamura G, Ikehara Y. 1986. Novel blockade by brefeldin A of intracellular transport of secretory proteins in cultured rat hepatocytes. J Biol Chem 261:11398–403.

136. Veillette M, Richard J, Pazgier M, Lewis GK, Parsons MS, Finzi A. 2016. Role of HIV-1 envelope glycoproteins conformation and accessory proteins on ADCC responses. Curr HIV Res 14:9–23.

137. Geleziunas R, Bour S, Wainberg MA. 1994. Cell surface down-modulation of CD4 after infection by HIV-1. FASEB J 8:593–600.

138. Levesque K, Zhao YS, Cohen EA. 2003. Vpu exerts a positive effect on HIV-1 infectivity by down-modulating CD4 receptor molecules at the surface of HIV-1-producing cells. J Biol Chem 278:28346–53.

139. Lama J, Mangasarian A, Trono D. 1999. Cell-surface expression of CD4 reduces HIV-1 infectivity by blocking Env incorporation in a Nef- and Vpu-inhibitable manner. Curr Biol 9:622–31.

140. Derdeyn CA, Decker JM, Sfakianos JN, Wu X, O’Brien WA, Ratner L, Kappes JC, Shaw GM, Hunter E. 2000. Sensitivity of human immunodeficiency virus type 1 to the fusion inhibitor T-20 is modulated by coreceptor specificity defined by the V3 loop of gp120. J Virol 74:8358–67.

141. Traunecker A, Schneider J, Kiefer H, Karjalainen K. 1989. Highly efficient neutralization of HIV with recombinant CD4-immunoglobulin molecules. Nature 339:68–70.

142. Byrn RA, Mordenti J, Lucas C, Smith D, Marsters SA, Johnson JS, Cossum P, Chamow SM, Wurm FM, Gregory T, Groopman JE, Capon DJ. 1990. Biological properties of a CD4 immunoadhesin. Nature 344:667–70.

143. Capon DJ, Chamow SM, Mordenti J, Marsters SA, Gregory T, Mitsuya H, Byrn RA, Lucas C, Wurm FM, Groopman JE, et al. 1989. Designing CD4 immunoadhesins for AIDS therapy. Nature 337:525–31.

144. Brand D, Srinivasan K, Sodroski J. 1995. Determinants of human immunodeficiency virus type 1 entry in the CDR2 loop of the CD4 glycoprotein. J Virol 69:166–71.

145. Adachi A, Gendelman HE, Koenig S, Folks T, Willey R, Rabson A, Martin MA. 1986. Production of acquired immunodeficiency syndrome-associated retrovirus in human and nonhuman cells transfected with an infectious molecular clone. J Virol 59:284–91.

146. Wang Q, Esnault F, Zhao M, Chiu TJ, Smith AB III, Nguyen HT, Sodroski JG. 2022. Global increases in human immunodeficiency virus neutralization sensitivity due to alterations in the membrane-proximal external region of the envelope glycoprotein can be minimized by distant State 1-stabilizing changes. J Virol 96:e0187821.

